# Intercellular transmission of a bacterial cytotoxic prion-like protein in mammalian cells

**DOI:** 10.1101/821215

**Authors:** Aida Revilla-García, Cristina Fernández, María Moreno-del Álamo, Vivian de los Ríos, Ina M. Vorberg, Rafael Giraldo

## Abstract

RepA is a bacterial protein that builds intracellular amyloid oligomers acting as inhibitory complexes of plasmid DNA replication. When carrying a mutation enhancing its amyloidogenesis (A31V), the N-terminal domain (WH1) generates cytosolic amyloid particles that are inheritable within a bacterial lineage. Such amyloids trigger in bacteria a lethal cascade reminiscent to mitochondria impairment in human cells affected by neurodegeneration. To fulfil all the features of a prion-like protein, horizontal (intercellular) transmissibility remains to be demonstrated for RepA-WH1. Since this is experimentally intractable in bacteria, here we transiently expressed in a murine neuroblastoma cell line the soluble, barely cytotoxic RepA-WH1(WT) and assayed its response to co-incubation with *in vitro* assembled RepA-WH1(A31V) amyloid fibres. In parallel, cells releasing RepA-WH1(A31V) aggregates were co-cultured with human neuroblastoma cells expressing RepA-WH1(WT). Both the assembled fibres and the extracellular RepA-WH1(A31V) aggregates induce, in the cytosol of recipient cells, the formation of cytotoxic amyloid particles. Mass spectrometry analyses of the proteomes of both types of injured cells point to alterations in mitochondria, protein quality triage, signalling and intracellular traffic.

**Summary blurb:** The horizontal, cell-to-cell spread of a bacterial prion-like protein is shown for the first time in mammalian cells. Amyloid cross-aggregation of distinct variants, and their associated toxicities, follow the same trend found in bacteria, underlining the universality of prion biology.

## Introduction

A hallmark of protein aggregates accumulating in human neurodegeneration is their ability to template the conformational change of soluble native protein molecules into highly ordered polymers with a cross-β structure termed amyloid (Eisenberg and Sawaya, 2017; Chiti and Dobson, 2017). This has been extensively characterized for the PrP prion in spongiform proteinopathies (Prusiner, 1998; Aguzzi and Falsig, 2012), the Aβ peptides (Eisele et al., 2010; Stöhr et al., 2012; Watts et al., 2014) and Tau (Frost et al., 2009; Clavaguera et al., 2009; Kfoury et al., 2012; Goedert et al., 2017) in Alzheimer’s and related dementias, or α-synuclein in Parkinson’s disease (Kordower et al., 2008; Luk et al., 2009; Luk et al., 2012; Prusiner et al., 2015). Protein aggregates are cell-to-cell transmissible through either tunnelling nanotubes, secretion as naked aggregates or packed into extracellular vesicles. These mechanisms underlie the stereotypical spreading of protein aggregates throughout connected brain regions (Polymenidou and Cleveland, 2012; Soto, 2012; Prusiner, 2013; Goedert, 2015; Jücker and Walker, 2018). Such ‘horizontal’ transmissibility, which partially resembles the infectivity of the PrP prion, qualifies Aβ peptides, Tau and α-synuclein as prion-like proteins (or prionoids) (Aguzzi and Lakkaraju, 2016; Scheckel and Aguzzi, 2018).

Intercellular aggregate transmission is not solely a characteristic of disease-associated proteins. Several proteins of lower eukaryotes have the ability to fold into self-templating protein aggregates that faithfully propagate in progeny and transmit to other cells during mating (Wickner et al., 2015). Interestingly, the prion-like behaviour is not restricted to the original host. For example, cell-to-cell propagation of amyloid aggregates has been successfully reported for the NM prion domain of the *Saccharomyces cerevisiae* translation termination factor Sup35 in mammalian cells (Krammer et al., 2009; Hofmann et al., 2013; Liu et al., 2016; Duernberger et al., 2018; Riemschoss et al., 2019). NM-Sup35 can also propagate in bacteria, provided that a specific prion-inducing amyloid required for the prionization of Sup35 in *S. cerevisiae* is also expressed in the recipient cells (Garrity et al., 2010). The other way around, both the amyloidogenic sequence stretch in RepA-WH1 (Gasset-Rosa and Giraldo, 2015) and the prion domain in bacterial CbRho (Yuan and Hochschild, 2017) can functionally replace Sup35 prionogenic sequences in a stop-codon read-through translation assay in yeast. However, the transmission of a bacterial prion or prion-like protein to mammalian cells and the assessment of its potential cytotoxicity have not been reported yet. This would complete the demonstration of the universality of the principles governing prion biology.

The bacterial prion-like protein RepA-WH1 is a synthetic model of amyloid disease built on RepA, a protein that controls plasmid replication through the assembly of amyloid oligomers that hamper premature rounds of origin firing (Giraldo et al., 2003; Molina-García et al., 2016). The WH1 domain in RepA forms stable dimers in solution. Upon allosteric binding to distinct ligands, dimers dissociate into metastable monomers that subsequently assemble as amyloid oligomers and fibres *in vitro* (Giraldo, 2007; Torreira et al., 2015). When expressed in *E. coli*, the hyper-amyloidogenic domain variant RepA-WH1(A31V) forms particles of various sizes, distributed across the bacterial cytosol (Fernández-Tresguerres et al., 2010). These particles are transmissible vertically (during cell division, i.e. from mother to daughter cells) as two distinct aggregate strains with remarkable appearances and phenotypes: elongated and mildly cytotoxic, and compact and acutely cytotoxic, respectively (Gasset-Rosa et al., 2014). Co-expression of soluble and aggregation-prone RepA-WH1 variants in *E. coli* demonstrated that the A31V variant can template its conformation on the parental WT protein (Molina-García and Giraldo, 2014). Systems analyses (Molina-García et al., 2017), together with *in vitro* reconstruction in cytomimetic lipid vesicles (Fernández et al., 2016b; Fernández and Giraldo, 2018), suggest that RepA-WH1(A31V) oligomers target the internal bacterial membrane, hampering proton motive force and thus ATP synthesis and transport through membranes, and enhance oxidative stress. In parallel, protein factors mounting the defence against stress and envelope damage co-aggregate with RepA-WH1(A31V) amyloids (Molina-García et al., 2017). Altogether, bacteria viability is severely compromised by RepA-WH1 amyloidosis, in a way resembling some of the central mitochondrial routes found in human amyloidoses (Haelterman et al., 2014; Norambuena et al., 2018; Mathys et al., 2019; Wang et al., 2019). However, *E. coli* is not suitable for addressing the issues of cell-to-cell transmissibility of protein aggregates and the subsequent intracellular amyloid cross-aggregation, since this Gram-negative bacterium does not uptake large protein particles due to the insurmountable obstacle of its three-layered cell envelope.

To explore the capability of the prion-like protein RepA-WH1 to propagate in a heterologous host, here we exposed murine neuroblastoma cells, transiently expressing mCherry-tagged soluble RepA-WH1(WT), to *in vitro* assembled RepA-WH1(A31V) amyloid fibres. In addition, we studied the intercellular induction of protein aggregation by co-culturing murine cells releasing RepA-WH1(A31V) aggregates with human neuroblastoma cells stably expressing soluble RepA-WH1(WT). Confocal microscopy and biochemical studies showed that the mammalian cells can take up RepA-WH1(A31V) amyloid fibres, that cross-seed the cytosolic aggregation of the endogenous RepA-WH1(WT) in the recipient cells. Moreover, co-culture of cells releasing the RepA-WH1(A31V) variant also induced the aggregation of RepA-WH1(WT) in bystander cells, suggesting intercellular transmission of RepA-WH1(A31V) seeds. In both setups of experimental transmission, the induced RepA-WH1(WT) aggregates were cytotoxic and amyloidogenic in nature, as indicated by cell viability assays and by their affinity for thioflavin-S, respectively. The observed toxicity was dependent on the expression of RepA-WH1(WT) in the recipient cells, suggesting that active assembly of protein aggregates and not simple uptake of toxic polymers was required. This reassures RepA-WH1, whose sequence shows no similarity matches in the human proteome, as a bio-safe prion-like protein. The analyses of the proteomes of either murine cells exposed to the *in vitro* assembled fibres, or human cells internalizing the extracellularly released aggregates, point to the impairment of mitochondria, intracellular trafficking and protein quality control networks in RepA-WH1 toxicity. The results presented here fulfil the validation of RepA-WH1 as a bacterial prion-like protein and reinforce the universality of the molecular and cellular basis of prion biology.

## Results

### Transient expression in mammalian cells of variants of the RepA-WH1 protein recapitulates their aggregation phenotypes in bacteria

Previous findings on the synthetic amyloid proteinopathy elicited in bacteria by the prion-like protein RepA-WH1 suggested that it might have features in common with a wide spectrum of human neurodegenerative amyloidoses. To assess if the model protein RepA-WH1, and its variants exhibiting distinct aggregation propensities in bacteria (Fernández-Tresguerres et al., 2010; Gasset-Rosa et al., 2014; Molina-Garcia et al., 2014), exhibit a comparable behaviour in mammalian cells, proteins were transiently expressed in murine N2a and human SH-SY5Y neuroblastoma cell lines, commonly used in studies about amyloid neurodegeneration, as well as in non-neuronal HeLa cells.

RepA-WH1 mutant variants were fused to the monomeric fluorescent protein mCherry in a constitutive expression plasmid (Figure S1A). These fusions, hereafter referred as WH1(WT/A31V/ΔN37)-mCherry for simplification, were the same that had been previously validated regarding their aggregation potential and toxicity in *E. coli* (Fernández-Tresguerres et al., 2010; Gasset-Rosa et al., 2014; Molina-Garcia et al., 2014; Molina-Garcia et al., 2017). While WH1(WT)-mCherry is soluble in the bacterial cytosol and non-cytotoxic, the hyper-amyloidogenic (A31V)-mCherry variant aggregates and is highly cytotoxic. WH1(ΔN37) is a deletion mutant lacking the amyloidogenic peptide stretch in RepA-WH1 that forms inclusion bodies. When this mutant is expressed in bacteria, it exhibits reduced toxicity compared to WH1(A31V)-mCherry. Cell lines were transfected with the plasmids coding for RepA-WH1 derivatives, or mCherry as a control. Soluble fractions of cell lysates were analyzed by Western blotting, 48 h after transient transfection, revealing variable protein expression in the three cell lines tested. The highest expression levels were observed in the N2a cells (Figure S1B). Variant WH1(ΔN37)-mCherry was not observed in any cell lysate. The N2a cell line was selected as an appropriate cell background for further exploring RepA-WH1 prion-like behaviour in mammalian cells. As previous work in bacteria had shown that WH1(ΔN37)-mCherry forms massive inclusion bodies (Gasset-Rosa et al., 2014; Molina-Garcia et al., 2014), we explored the presence of this variant in the insoluble lysate fraction of the N2a cells. WH1(ΔN37)-mCherry was clearly located in the pellet, according to Western blot analysis (Figure S1B, right panel), thereby confirming that this mutant also forms insoluble aggregates in the mammalian cytosol.

To evaluate the possible cytotoxicity of RepA-WH1, the N2a transfected cells were observed by means of confocal microscopy. As observed in Figure 1A, WH1(WT)-mCherry and the mCherry control remained phenotypically soluble (i.e., they showed diffuse cytoplasmic fluorescence) when overexpressed, which demonstrated that none of them has tendency towards spontaneous aggregation. On the contrary, for the WH1(A31V)-mCherry mutant, small red foci were clearly detected in the cytosol, which correlated with its natural propensity to aggregate in bacteria (Fernández-Tresguerres et al., 2010; Gasset-Rosa et al., 2014; Molina-Garcia et al., 2014). Finally, larger intracellular foci, distinct to those for WH1(A31V)-mCherry, were observed when WH1(ΔN37)-mCherry was expressed.

**Fig. 1.**
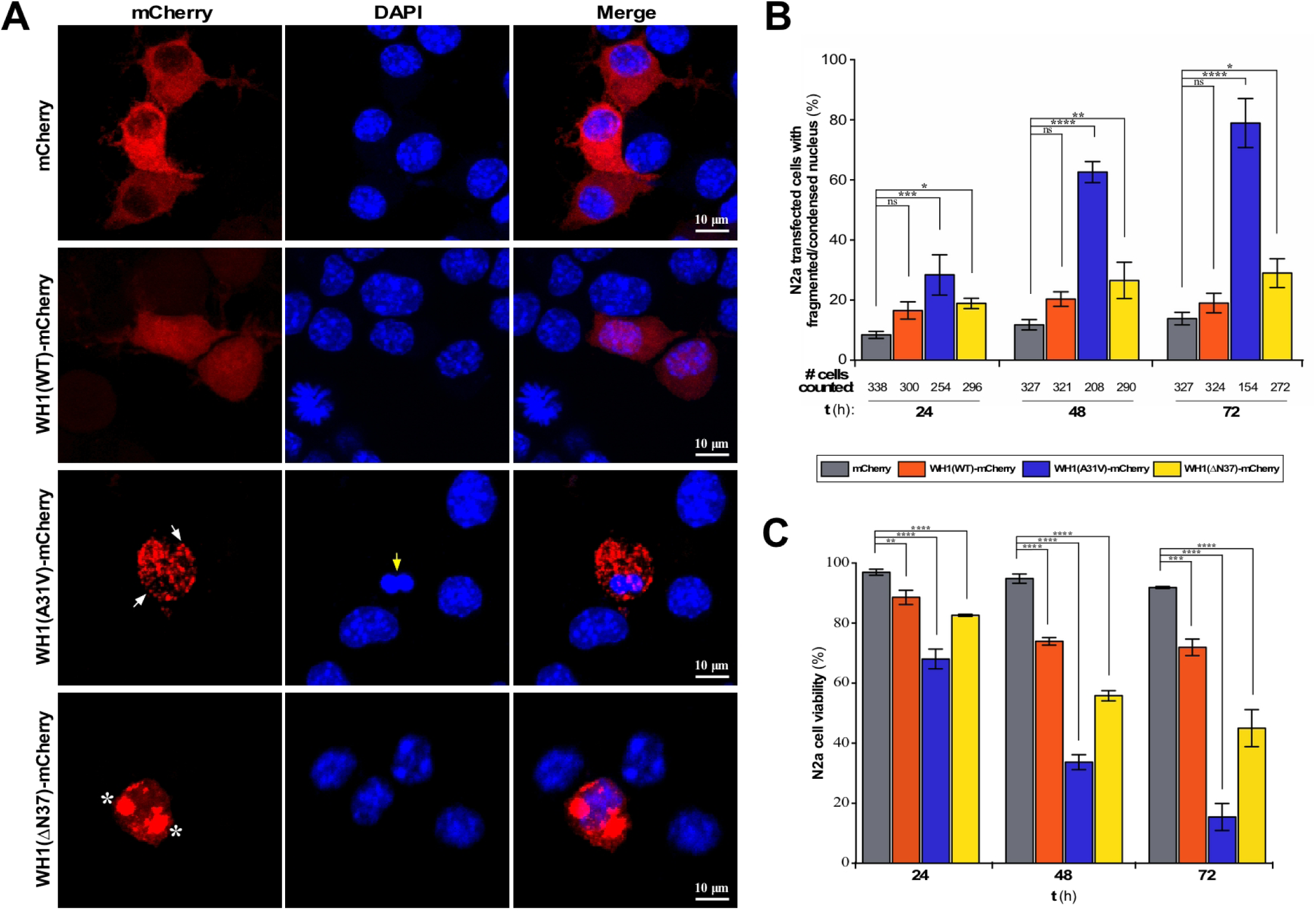
Expression of WH1(A31V)-mCherry in the cytosol of murine N2a cells. (**A**) Confocal maximum-intensity projection images of N2a cells expressing WH1(WT/A31V/ΔN37)-mCherry, or mCherry as a control, 48 h after being transiently transfected (red). White arrows mark the small WH1(A31V)-mCherry fluorescent foci, whereas the large WH1(ΔN37)-mCherry inclusions are indicated with asterisks. Nuclei were stained with DAPI (blue). Yellow arrow points to an altered nucleus. (**B**) Quantitative analysis of condensed/fragmented nuclei in N2a transfected cells that expressed for 24, 48 and 72 h either WH1(WT/A31V/ΔN37)-mCherry or the mCherry control. The total number of cells of each type expressing the mCherry fluorescence is displayed below the X-axis. (**C**) N2a cell viability, quantitatively estimated from a MTT reduction assay, in response to transient expression of the RepA-WH1 variants for 24, 48 and 72 h. The inferred cytotoxicity of the proteins expressed was: WH1(A31V) ≫ WH1(ΔN37) > WH1(WT) > mCherry. In (B) and (C), bars correspond to the mean values from three independent transfections (n = 3). One-way ANOVA test was performed (*, p < 0.1; **, p < 0.01; ***, p < 0.001; ****, p < 0.0001; ns: not significant).

Remarkably, the inspection of nuclei morphology by confocal microscopy revealed the noticeable presence of N2a cells exhibiting a fragmented or condensed nucleus upon WH1(A31V)-mCherry expression, a sign for cell death. This phenotype was rare in cells expressing WH1(WT/ΔN37)-mCherry, or the mCherry control (Figure 1A). Quantitative analysis of the cells showing nuclear condensation/fragmentation at 24, 48 and 72 h after transfection revealed that such difference was statistically significant (Figure 1B). A substantial decrease in the number of WH1(A31V)-mCherry expressing cells was detected at 48 h and, specially, 72 h after transfection, while similar numbers of either mCherry control or WH1(WT/ΔN37)-mCherry transfected N2a cells were counted over time. Such reverse correlation between fragmented/condensed nuclei and cell count supports the cytotoxicity of WH1(A31V).

To further assess the cytotoxic effect of WH1(A31V)-mCherry overexpression in N2a cells, cell viability was estimated by monitoring the activity of NAD(P)H-dependent oxidoreductases 24, 48 and 72 h after transfection (Figure 1C). The MTT reduction assay revealed a decrease in cell viability, compared to the mCherry control, for cells expressing either WH1(A31V)-mCherry or WH1(ΔN37)-mCherry. This toxicity was time-dependent and most pronounced for cells expressing WH1(A31V)-mCherry, resulting in a decrease of cell viability by about 60% at 48 h, to reach 90% at 72 h. On the contrary, no time-dependent changes in cell viability were detected for mCherry and WH1(WT)-mCherry.

To verify that the cytotoxic aggregates formed by RepA-WH1 in the N2a cells were of amyloid nature, we relied on thioflavin-S (ThS), an amyloidotropic fluorophore that stains RepA-WH1 amyloids in bacteria (Fernández-Tresguerres et al., 2010; Gasset-Rosa et al., 2014; Giraldo, 2019). Forty-eight hours after WH1(WT/A31V/ΔN37)-mCherry or mCherry transfection, staining with ThS was performed and cells were visualized by confocal microscopy (Figure 2). Image analysis revealed that in the N2a cells expressing WH1(A31V)-mCherry, the aggregated foci were strongly stained by ThS, an indication of their amyloid nature. In contrast, weak, near background ThS staining was observed for cells expressing either the mCherry control, the soluble WH1(WT)-mCherry protein, or for the large WH1(ΔN37)-mCherry aggregates. Reduced binding of the fluorophore was previously considered indicative of the amorphous aggregation of WH1(ΔN37)-mCherry as bacterial inclusion bodies (Gasset-Rosa et al., 2014).

**Fig. 2.**
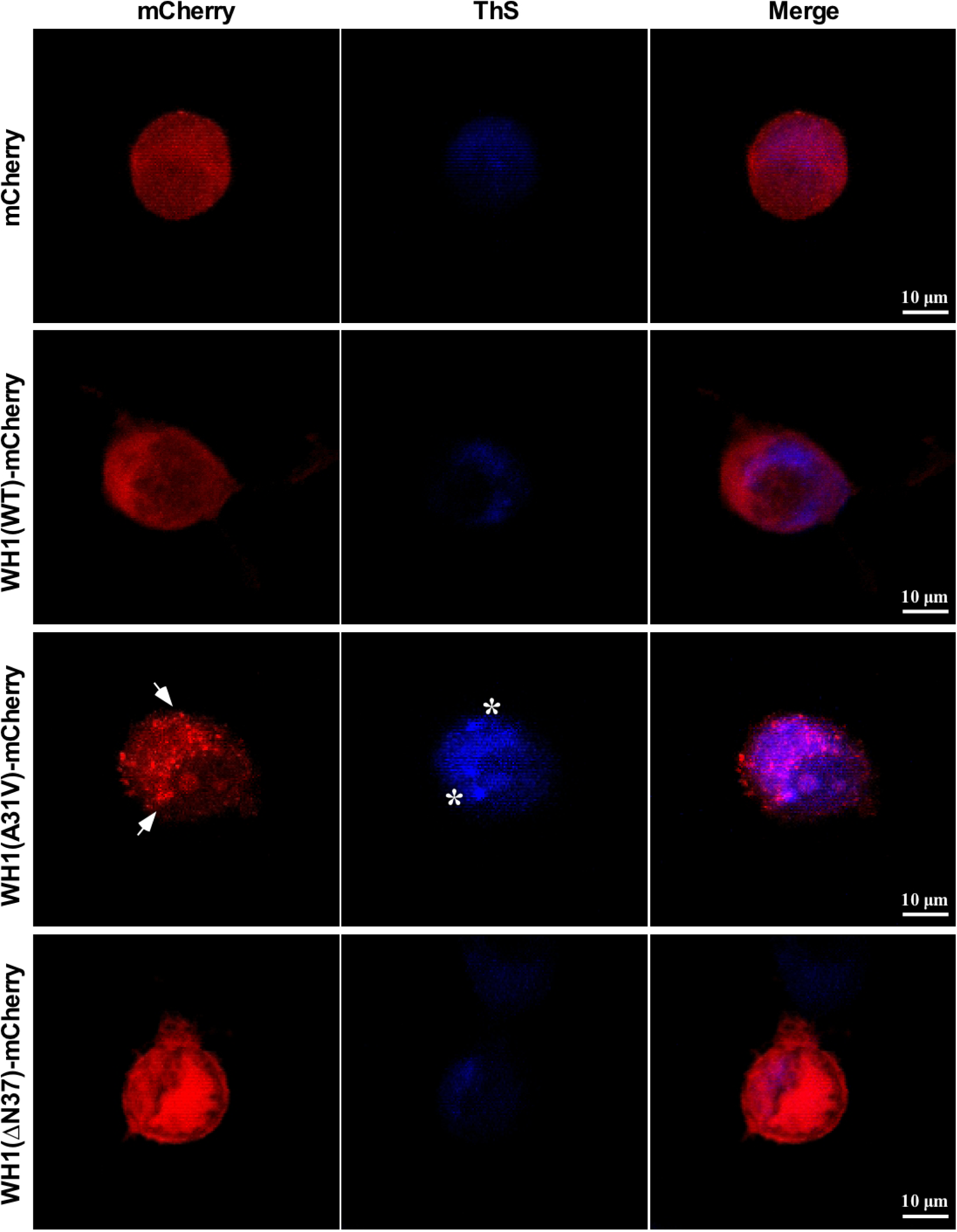
The particles assembled by WH1(A31V)-mCherry at the cytosol of murine N2a cells are amyloids. Confocal maximum-intensity projection images of N2a cells transiently expressing for 48 h WH1(WT/A31V/ΔN37)-mCherry, or mCherry as control (red). Cells were stained with ThS (blue) for amyloid detection. Arrows mark examples of the intracellular WH1(A31V)-mCherry puncta, whereas some of the ThS-positive foci are indicated with asterisks. Just the WH1(A31V)-mCherry particles are amyloid by nature.

### *In vitro*-assembled RepA-WH1(A31V) amyloid fibres co-localize with RepA-WH1(WT) transiently expressed in the cytosol of murine cells

One of the most remarkable properties of amyloid aggregates is their inherent self-propagation potential, being able to template the amyloid conformation on soluble molecules of the same protein. For RepA-WH1 this had been shown in *E. coli in vivo* by co-expressing WH1(A31V) and WH1(WT) labelled with different fluorescent proteins (Molina-García and Giraldo, 2014). Thereby, validation of RepA-WH1 as a ‘generic’ model for amyloidosis required testing its ability to cross-seed aggregation within mammalian cells.

We explored whether exogenous, *in vitro* assembled WH1(A31V) fibres (Figure S2), whose amyloid nature has been well established (Giraldo, 2007; Fernández-Tresguerres et al., 2010; Moreno del Álamo et al., 2015; Torreira et al., 2015; Fernández et al., 2016a), could trigger the aggregation of the otherwise soluble WH1(WT) protein (Figure 1A, second row) in the cytosol of mammalian cells. To this aim, N2a cells transiently expressing WH1(WT)-mCherry in the cytosol were exposed to *in vitro* pre-assembled WHI(A31V)-Alexa 488 labelled amyloid fibres (for simplification, hereafter WH1(A31V) fibres) (Figure 3A). These were mechanically fragmented before incubation with cells to get fibre sizes more suitable for their uptake (see Materials and Methods) (Figure S2). To check for the specific requirement of the WH1(WT) domain as the target for aggregation templating by WH1(A31V) fibres, control cells expressing just mCherry were incubated with the fibres in parallel. Confocal microscopy sections revealed that the WH1(A31V) fibril particles were efficiently internalized by the N2a cells expressing either mCherry or WH1(WT)-mCherry (Figure 3B). In the latter, fibrils co-localized with the endogenous WH1(WT)-mCherry in small, dot-like cytosolic aggregates (Figure 3B, arrows). Approximately 70% of the WH1(WT)-mCherry expressing cells displayed red puncta upon incubation with the fibres, while spontaneous foci formation of WH1(WT)-mCherry was detected in < 35% of the cells after 48 h (Figure 3C). These figures significantly differed from < 20% of foci-bearing cells counted for the mCherry control, irrespective of the presence or absence of added fibrils (Figure 3C). Overall, these observations support that induction of foci by WH1(A31V) amyloid fibres requires expression of a homologous protein substrate in the recipient cells.

**Fig. 3.**
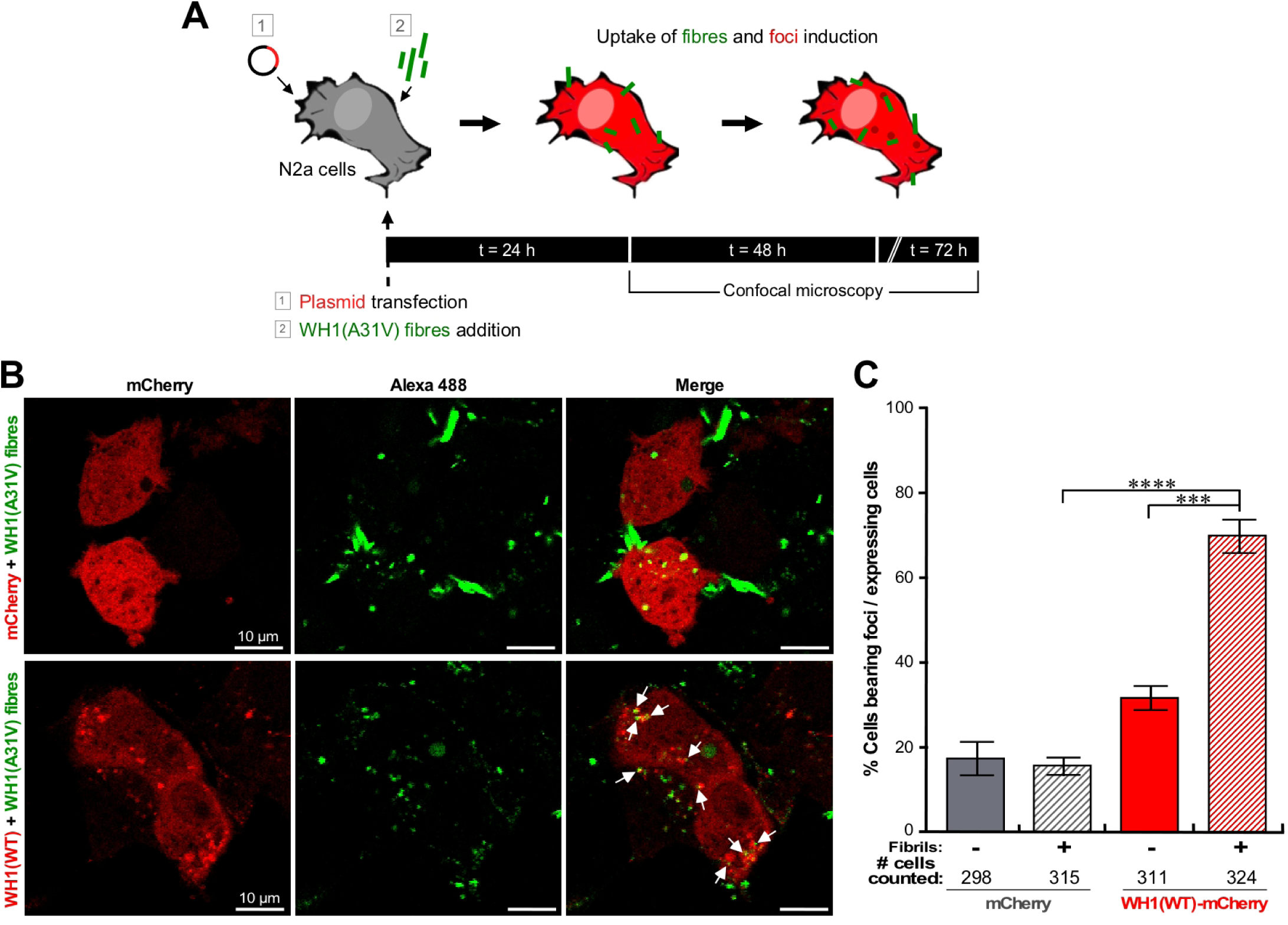
Uptake of *in vitro*-assembled, Alexa 488-labelled WH1(A31V) amyloid fibres by murine N2a cells results in the formation of foci by endogenous WH1(WT)-mCherry. (**A**) Experimental setup to study the induction of endogenous WH1(WT)-mCherry foci upon the uptake of exogenous WH1(A31V) fibres. N2a cells were transiently transfected with plasmids expressing WH1(WT)-mCherry or the mCherry control (1) and immediately incubated with *in vitro*-assembled WH1(A31V) fibrils (2) for 24, 48 and 72 h. (**B**) Confocal section images of the experiment outlined in (A) (48 h after fibre addition, unfixed cells). Exogenous WH1(A31V) fibres (green) were taken up by N2a cells expressing (red) either mCherry (top), or WH1(WT)-mCherry (bottom). The generated intracellular WH1(WT)-mCherry amyloid foci are indicated by arrows. (**C**) Percentage of mCherry or WH1(WT)-mCherry expressing N2a cells bearing fluorescent foci 48 h after incubation with WH1(A31V) fibres. Bars in the graph display the means from three independent experiments (n = 3). The total number of cells of each type counted is displayed below the X-axis. For statistical analysis, Student’s *t*-test was performed (***, p < 0.001; ****, p < 0.0001).

As a consequence of amyloid templating, it can be assumed that the foci induced in mammalian cells by the internalized amyloid fibres are amyloid aggregates. In order to qualitatively assess this point, ThS staining was performed (Figure 4). Confocal microscopy images obtained after 48 h of fibre addition to the cell cultures revealed the selective binding of ThS to the induced cytosolic WH1(WT)-mCherry aggregates. On the contrary, no intense ThS staining was observed for cells expressing mCherry, attributable to background fluorescence. These findings are compatible with the results of ThS binding to the intracellular aggregates generated upon transient expression of WH1(A31V)-mCherry (Figure 2). Therefore, ThS staining provides evidence for the amyloid nature of the WH1(WT)-mCherry aggregates as templated by the WH1(A31V) fibres in the cytoplasm of murine N2a cells.

**Fig. 4.**
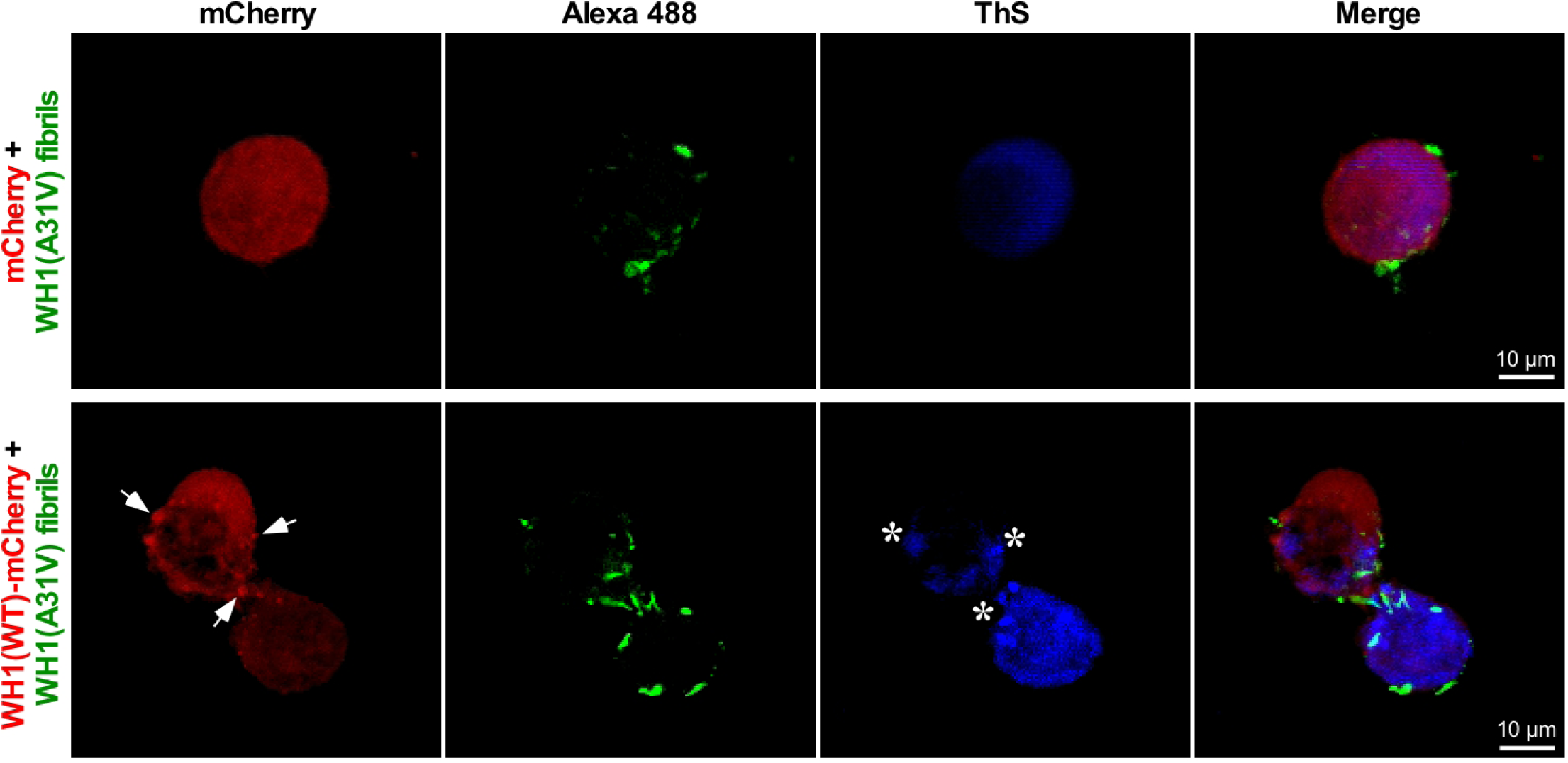
The particles induced by transfected WH1(A31V) fibres on WH1(WT)-mCherry at the cytosol of murine N2a cells are amyloidogenic. Confocal maximum-intensity projections of N2a cells expressing soluble mCherry or WH1(WT)-mCherry (red), incubated for 48 h with *in vitro*-assembled, Alexa 488-labelled WH1(A31V) fibres (green). Cells were then stained with ThS (blue) for amyloid detection. Three of the fibre-nucleated WH1(WT)-mCherry puncta (arrows) that were also positive for ThS (asterisks) are highlighted.

### RepA-WH1(A31V) released from murine cells can cross-seed amyloid aggregation in the cytosol of recipient human cells stably expressing RepA-WH1(WT)

The studies described above provided evidence for the ability of WH1(A31V) to cross-seed WH1(WT) amyloidogenesis in the cytosol of mammalian cells. To further explore this, we aimed to generate stable donor cell lines in which the WH1 variants and controls, fused to the green fluorescent protein TagGFP2, could be expressed under an inducible promoter (pTRE3G Tet-ON vector, genome-integrated THN-rtTA transactivator; see Materials and Methods) (Figure S3A). We chose SH-SY5Y as the parental donor cell line as transgene expression in that background was less pronounced than in N2a cells (Figure S1B). Unfortunately, we were unable to get any clone expressing WH1(A31V)-TagGFP2, suggesting that, even under a tightly regulated gene expression, any transcriptional leak of this mutant variant results in acute cytotoxicity. For the WH1(WT)-TagGFP2 and the control TagGFP2 constructs, efficient doxycycline (Dox)-induced expression was verified through Western blot (Figure S3B) and flow cytometry (Figure S3C) analyses. Confocal microscopy imaging indicated that both the THN-rtTA/WH1(WT)-TagGFP2 and THN-rtTA/TagGFP2 double-stable clones showed an homogeneous, diffuse expression throughout the entire cytosol (Figure S3D).

The horizontal, cell-to-cell spreading of RepA-WH1, a key feature to qualify it as a prion-like protein (Aguzzi and Lakkaraju, 2016), was then explored. To this aim, murine N2a donor cells transiently expressing either WH1(A31V)-mCherry or WH1(WT)-mCherry (Figure 1A) were co-cultured with THN-rtTA/WH1(WT)-TagGFP2 recipient cells, which stably expressed soluble WH1(WT)-TagGFP2 upon Dox addition (Figure S3D), for a period of up to a week (Figure 5A). To control for non-specific spontaneous aggregation in the human cells, recipient cultures not mixed with donors were similarly processed in parallel. The formation of foci in the recipient cells was assessed by confocal microscopy imaging every 24 h (Figure 5B,C, left). Noticeably, small green fluorescent puncta were detected over time in the THN-rtTA/WH1(WT)-TagGFP2 recipient cells when the donor N2a cells expressing the hyper-amyloidogenic variant WH1(A31V)-mCherry were added, peaking at 5 days (Figure 5C, left-bottom) and decreasing thereafter, which suggests killing of the cells. On the contrary, no substantial formation of WH1(WT)-TagGFP2 foci was observed upon addition of the N2a donors transiently expressing WH1(WT)-mCherry (Figure 5C, left-top) to the human recipient cells.

**Fig. 5.**
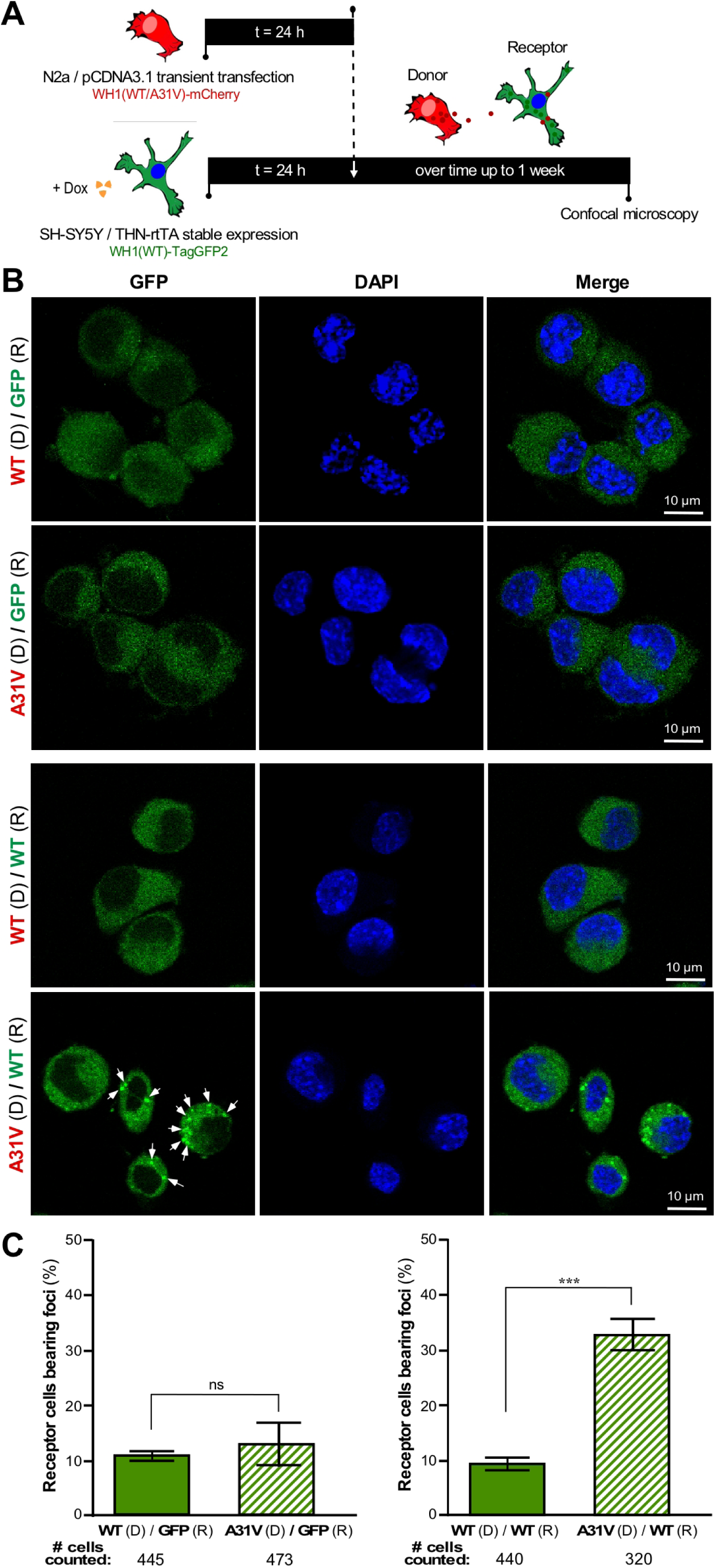
Uptake by human SH-SY5Y cells of WH1(A31V)-mCherry generated by murine N2a cells results in formation of foci by endogenously expressed WH1(WT)-TagGFP2. (**A**) Experimental setup to study horizontal propagation of WH1(A31V) between cell lines. Twenty-four h after being transfected with plasmids, donor N2a cells transiently expressing soluble WH1(WT)-mCherry, or the amyloidogenic WH1(A31V)-mCherry, were co-cultured for up to 7 days with THN-rtTA/SH-SY5Y receptor cells stably expressing, upon Dox induction, either WH1(WT)-TagGFP2 or TagGFP2. (**B**) *Left*: Confocal sections of cells in cultures as sketched in (A) in which the receptor cells (R) were expressing the TagGFP2 control, while the donor cells (D) were either WH1(WT)-mCherry or WH1(A31V)-mCherry. No large fluorescent puncta were observed after 5 days. DAPI staining of nuclear DNA was carried out. *Right*: Percentage of THN-rtTA/SH-SY5Y/TagGFP2 receptor cells (R) bearing intracellular foci after 5 days of co-culture with transiently transfected WH1(WT)-mCherry or WH1(A31V)-mCherry N2a donors (D). (**C**)*Left*: Confocal sections of cells in cultures as those in (B), but in which the receptors (R) were expressing WH1(WT)-TagGFP2 instead of just TagGFP2. Bright green spots (arrows) were differentially promoted by the cells donating WH1(A31V)-mCherry, but not by those expressing WH1(WT)-mCherry. *Right*: Quantitative analysis of intercellular propagation of RepA-WH1. Percentage of THN-rtTA/SH-SY5Y/WH1(WT)-TagGFP2 receptor cells (R) bearing intracellular foci after 5 days of co-culture with transiently transfected WH1(WT)-mCherry or WH1(A31V)-mCherry N2a donors (D). Bars displayed in both histograms are the mean of three independent experiments (n = 3). The total number of cells of each type counted is displayed below the X-axes. For statistical analysis, Student’s *t*-test was performed (***, p < 0.001; ns, not statistically significant).

Fluorescent puncta were not apparent either for their control counterparts, in which donors expressing WH1(A31V/WT)-mCherry were added to cultures of recipient cells THN-rtTA/TagGFP2 (Figure 5B, left). These results fully parallel (and strengthen) the observations made when exogenously added, *in vitro*-assembled WH1(A31V) fibres induced the formation of WH1(WT)-mCherry foci in the N2a cells (see above; Figure 3). Five days co-cultures were selected for quantitative image analysis of the fraction of cells that exhibited green fluorescent foci. Just 10-12% of the human control cells endogenously expressing TagGFP2 showed some fluorescent spots, irrespective of being co-cultured with murine cells expressing WH1(A31V)-mCherry or WH1(WT)-mCherry. Puncta were apparent in ≈35% of the recipient cells expressing WH1(WT)-TagGFP2 only if the hyper-amyloidogenic WH1(A31V)-mCherry variant was expressed by the donor murine cells (Figure 5B,C, right).

While WH1(A31V)-mCherry aggregates must be first released (either through exocytosis or, more likely, upon cell death) to the culture medium from the donor cells and subsequently gain access into the recipient cells, it is noteworthy that red fluorescent WH1(A31V)-mCherry particles were rarely detected at the intercellular space. Although it is possible that the loss of such particles may occur during the manipulation of the samples (e.g., for cell passages or microscopy), the most likely explanation is that their size range is beyond the resolution of conventional confocal imaging (Hofmann et al., 2013; Liu et al., 2016).

In conclusion, these results showed significant differences in the intercellular ‘infectivity’ of soluble WH1(WT)-mCherry and hyper-amyloidogenic WH1(A31V)-mCherry. Cells suffering from the RepA-WH1(A31V) amyloidosis were highly proficient in the propagation of an aggregation-related phenotype, which was consistent with the results obtained for the *in vitro* pre-assembled RepA-WH1(A31V) fibrils. Altogether, these parallel observations suggest that the bacterial RepA-WH1 protein indeed follows in mammalian cells a prion-like mechanism for horizontal transmission.

### A proteomic outline of the cytotoxicity of RepA-WH1(A31V) amyloids in mammalian cells

In a previous study, we characterized the systems-wide response of *E. coli* to WH1(A31V)-mCherry expression to identify pathways potentially affected by the bacterial amyloidosis (Molina-García et al., 2017). To assess cellular responses to the toxic WH1(A31V) amyloids, we performed parallel proteomic studies on three groups of samples: i) N2a cells transiently expressing WH1(WT)-mCherry or mCherry, collected after 48 h of incubation with WH1(A31V) fibres (see Figure 3; Supplemental Dataset 1); ii) THN-rtTA/SH-SY5Y cells expressing WH1(WT)-TagGFP2 or TagGFP2, harvested after 5 days of induction with Dox in co-culture with N2a cells expressing WH1(A31V)-mCherry (see Figure 5; Supplemental Dataset 2); and iii) control naïve N2a or THN-rtTA/SH-SY5Y cells incubated, respectively, with WH1(A31V) fibres or WH1(A31V)-mCherry donor cells.

The subsets of the proteomes that were exclusively found in recipient cells expressing WH1(WT)-mCherry (N2a cells, mice) or WH1(WT)-TagGFP2 (THN-rtTA/SH-SY5Y cells, human), and not in cells just expressing their respective fluorescent tag controls, upon fibre-induced or cell-to-cell spread of amyloidogenesis reflect a broader response for latter experimental scenario (Figures 6 and S4). Albeit the individual proteins identified are not the same (perhaps also due to host species-specific features), both datasets point to shared pathways affected in the RepA-WH1 amyloidosis. These include distinct subcellular organelles, specially mitochondria and the endoplasmic reticulum, and processes, among which metabolism and signalling are the most frequent hits. Cluster analyses further refine these findings, confirming that mitochondria are at the very core of the RepA-WH1 amyloidosis under both experimental approaches (Figure S4), with Complex I of the respiratory chain (NADH:ubiquinone oxidoreductase) as the most prominent node. In addition, protein quality control machinery (the proteasome, ubiquitin-ligases and chaperones), cell signalling and intracellular trafficking networks, linked to protein secretion, are affected by WH1(A31V) uptake.

**Fig. 6.**
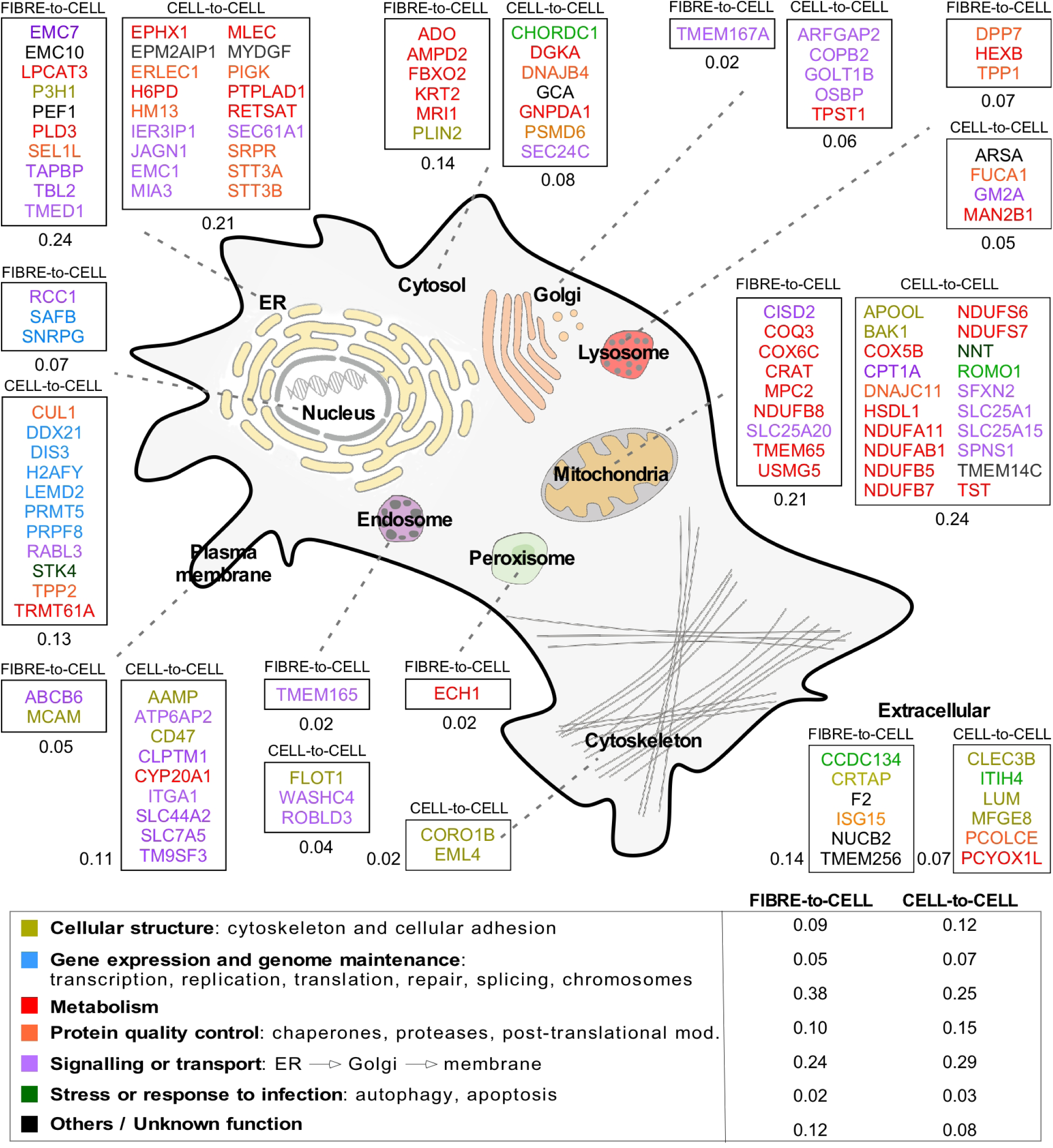
Proteome-wide responses of mammalian cells expressing WH1(WT) to either *in vitro* assembled WH1(A31V) fibres or WH1(A31V)-mCherry released from co-cultured cells. The cartoon displays, overlaid to the sketch of a mammalian cell, the names of the proteins differentially and uniquely found in cells expressing WH1(WT)-mCherry, and on which amyloids were templated by either fibre-to-cell or cell-to-cell transmission. Proteins are grouped in boxes according to their subcellular location, with numbers representing the fraction of the proteome dataset allocated to each compartment. Colour coding refers to the main function reported in the literature for these proteins (see the legend below, including the fraction of the proteins placed in each functional group).

Interestingly, similarities between the response of mammalian mitochondria to amyloid-promoted neurodegeneration and how *E. coli* reacts to RepA-WH1 amyloidosis were previously proposed on the basis of systems analyses carried out in the latter host (Molina-García et al., 2017). The results presented here reinforce this view and make sense in light of the common ancestry of Gram-negative bacteria and mitochondria, validating RepA-WH1 as a minimal, autonomous proxy to sketch core mitochondrial routes involved in human amyloidoses.

## Discussion

The ability of highly ordered protein aggregates to intercellulary template their amyloid structure on soluble molecules of the same protein found in naïve recipient cells has recently emerged as a hallmark of human neurodegenerative diseases. In common with the infectious propagation of the PrP prion, cell-to-cell transmission of prion-like protein aggregates has been established for α-synuclein, Tau, Aβ peptides, and it is now being explored for many other proteins involved in neurodegeneration (Polymenidou and Cleveland, 2012; Soto, 2012; Prusiner, 2013; Goedert, 2015; Jücker and Walker, 2018). The proteins involved in such aggregation cross-seeding *in vivo* are thus considered prion-like proteins or prionoids (Aguzzi and Lakkaraju, 2016; Scheckel and Aguzzi, 2018). In these cases, the intercellular traffic is likely mediated by generation of extracellular vesicles from the donor cells and subsequent endocytic uptake by the recipient cells (Liu et al., 2016).

RepA-WH1 is a bacterial prion-like protein that recapitulates in *E. coli* many of the essential features of an amyloid proteinopathy, including the formation of cytotoxic amyloid aggregates (Fernández-Tresguerres et al., 2010; Molina-García et al., 2017), the existence of strains that give rise to distinct phenotypes, and their epigenetic vertical (mother-to-daughter cells) transmission coupled to cell division (Gasset-Rosa et al., 2014). However, its ability to be horizontally (cell-to-cell) transmitted as a prion-like protein had not been addressed yet in bacteria, due to the stringent barrier that to the uptake of large protein aggregates impose the peptidoglycan and, in Gram-negative bacteria, also the outer membrane. We have now used mammalian cultured cells and fluorescent-labelled proteins to address the issue of the horizontal transmissibility of RepA-WH1. We first transiently expressed distinct variants of RepA-WH1 in murine cells, which led to the formation of amyloids with similar degrees of cytotoxicity to those described in bacteria, i.e., WH1(A31V) ≫ WH1(ΔN37) > WH1(WT) (Gasset-Rosa et al., 2014; Molina-García et al., 2017). Then incubation of those cells with *in vitro*-assembled amyloid WH1(A31V) fibres showed their ability to induce the formation of foci by WH1(WT), which is otherwise soluble as expressed in the cytoplasm of the recipient cells. In a subsequent step, we generated a stable and inducible cell line expressing soluble WH1(WT), which assembled as fluorescent puncta upon exposure to WH1(A31V) particles generated by donor cells. The results from these experiments fulfil the last stalling requirement to qualify RepA-WH1 as a prion-like protein: its ability to be horizontally transmissible.

It is noteworthy that, under our experimental conditions, the uptake of recombinant WH1(A31V) amyloids by cells not expressing WH1(WT), but just the fluorescent proteins to which the prion-like protein was tagged, does not result in any evident toxicity for the recipient cells. Although there is no protein with a noticeable degree of sequence identity with RepA-WH1 in the human proteome (BLAST best match E-value = 0.11), it has been recently suggested that bacterial extracellular amyloids may template amyloidogenesis on human proteins implicated in neurodegeneration in spite of the lack of significant sequence similarities (Friedland and Chapman, 2017; Perov et al., 2019). This could constitute a matter of concern on the biosafety of bacterial amyloids. In contrast, the inability of WH1(A31V) amyloids to induce either intracellular aggregation or toxicity in naïve mammalian cells not expressing RepA-WH1 qualifies this protein as biosafe.

In the context of the experiment in which amyloidogenesis was induced by adding exogenous fibres, it is most relevant that the intracellular foci in the recipient cells were templated by a well characterized amyloid structure assembled *in vitro* (Giraldo, 2007; Torreira et al., 2015), which in turn was seeded by intracellular aggregates extracted from *E. coli* cells experiencing WH1(A31V) amyloid toxicity (Fernández-Tresguerres et al., 2010). Therefore, it can be assumed (Luk et al., 2009; Stöhr et al., 2012; Lu et al., 2013; Frederick et al., 2014; Qiang et al., 2017) that the intracellular amyloid (ThS-reactive) foci in the recipient murine cells may have the same intimate structure as the aggregates reported to kill bacterial cells.

The toxicity of the amyloid proteins involved in neurodegeneration can be due to the loss, on their aggregation, of an essential cellular function and/or to the gain of an emergent functionality by the aggregates (Chiti and Dobson, 2017), e.g., upon the assembly of oligomeric membrane pores (Fernández et al., 2016b). In the case of the phenotype caused in the mammalian cells by the expression of WH1(A31V) amyloids (Figures 1 and 2), or by their uptake and subsequent templating on WH1(WT) in the recipient cells (Figures 3–5), cytotoxicity is necessarily an intrinsic property of the aggregates themselves, because *a priori* no function is associated with this bacterial prion-like protein in the heterologous mammalian host. Very recently, RepA-like proteins with a conserved WH1 domain (Kilic et al., 2019) have been described as nearly the only gene products encoded by small mobile genetic elements that were isolated in dairies and meat (BMMFs), with a probable origin in cattle microbiota, but also in samples from patients with cancer and multiple sclerosis (de Villiers et al., 2019). These findings open the intriguing possibility that the cytotoxicity described here for RepA-WH1 might have broader relevance in human disease.

Mass spectrometry analyses of the proteins specifically detected upon the uptake of RepA-WH1(A31V) under the two main scenarios of experimental transmission addressed (Figure 6) provided a picture sharing some similarities to cells in neurodegeneration. In the case of exogenous nucleation of WH1(WT) amyloids by WH1(A31V) fibres, proteins annotated as involved in neurodegenerative diseases were identified (Crat, Hexb, Pld3 and Sel1l), while others were related to ubiquitin ligases and proteasome-mediated degradation pathways (Fbxo2 and Isg15). Upon the uptake of secreted/released WH1(A31V) particles, proteins linked to neurodegeneration (Arsa, Epm2aip1 and Hm13) also showed-up, together with co-chaperones (Chordc1, Dnajb4 and Dnajc11) and proteins involved in vesicular trafficking (Arfgap2, Copb2, Dis3 and Flot1) or in apoptosis (Bak, Cul1, Ier3ip1, Mfge8, Psmd6, Tpp2 and Stk4). It is noteworthy that the latter participates in chromatin condensation, as observed in Figure 1B. Mitochondrial proteins as well as those located at the endoplasmic reticulum were the most common hits, preferentially proteins directly related with bioenergetics (respiratory chain) and with secretory pathways, respectively. Two proteins are worth noting in the context of what is known on RepA-WH1(A31V) toxicity in *E. coli*: Apool, which binds to cardiolipin, the bacterial and mitochondrial-specific acidic phospholipid that promotes the formation of pores by the prionoid in model membranes (Fernández et al., 2016b), and Romo1, a protein that enhances the formation of reactive oxygen species (ROS) in mitochondria, usually as an anti-microbial response. Romo1 might be playing the same role, albeit through a different mechanism, that the auto-oxidation of the alternative dehydrogenase NdhII plays in bacteria, i.e., constituting the first step in the oxidative cascade leading to cell death (Molina-García et al., 2017). Attempts to counteract ROS generation in mitochondria seem to rely on Nnt, which generates NADPH and thus reduction power to detoxify free radicals. Interestingly, a central role in the cytotoxicity of the PrP prion has been proposed for NADPH oxidases (Sonati et al., 2013).

Several systems analyses, based on differential transcriptomics and proteomics, have been reported for conditions related to neurodegenerative amyloidoses (Hwang et al., 2009; Olzscha et al., 2011; Hosp et al., 2015; Woerner et al., 2016; Kim et al., 2016; Seyfried et al., 2017; Markmiller et al., 2018; Mathys et al., 2019). One-third of the proteins identified in our analysis (Figure 6) were also found among the genes overexpressed, or the proteins captured within aggregates, in those datasets, thus defining a subset of proteins (Table S1) distinctive of a ‘generic’ cellular model of amyloidosis, as it can be outlined with the synthetic bacterial prion-like RepA-WH1 protein.

## Materials and Methods

### Plasmid Constructions

#### Generation of the pcDNA3.1 vectors

The gene fusions *repA-WH1(WT)-mCherry, repA-WH1(A31V)-mCherry* and *repA-WH1(ΔN37)-mCherry*, or *mCherry* were amplified by PCR (5’-CCGGGTACCATGGGCAGCAGCCATCATC and 5’-CGGGAATTCTTACTTGTACAGCTCGTCCAT primers; Pfu DNA polymerase, Promega) from the pRG*rectac*-NHis constructions (Fernández-Tresguerres et al., 2010). The PCR products were cloned into pcDNA3.1 (Invitrogen) after cutting with KpnI (5’end) and EcoRI (3’end) (Fig. S1A). Constructs were screened by colony PCR and restriction enzyme digestion, and confirmed by Sanger sequencing (Secugen: https://www.secugen.es).

#### Generation of the pTRE3G vectors

For generation of SH-SY5Y stable cell lines expressing WH1(WT)-TagGFP2, or TagGFP2 as a control, the *TagGFP2* sequence of the pTagGFP2-N vector (Evrogen) was amplified via PCR (5’-ATAAGAATTCCGGAATGAGCGGGGGCGAGGAG and 5’-CGGGGAATTCCATATGTTACCTGTACAGCTC primers) and cloned into pcDNA3.1-mCherry (see above) via BspEI and EcoRI digestions, thus replacing the red by the green fluorescent protein. The RepA-WH1(WT) domain was then excised form pcDNA3.1-*WH1(WT)-mCherry* and fused in frame with TagGFP2 by digestion with SacII and BspEI.

For the Tet-ON inducible expression plasmid pTRE3G (Clontech), *WH1(WT)-TagGFP2* and *TagGFP2* genes were amplified by PCR (5’-GAAGATCTATGGGCAGCCATCATCATCA and 5’-CCGGAATTCCATATGTTACCTGTACAGCCTC primers). The amplified PCR fragments were subcloned into pTRE3G with BglII and NdeI to obtain pTRE3G-*TagGFP2* and pTRE3G-*WH1(WT)-TagGFP2* (Fig. S3A).

#### Generation of pET-*WH1(A31V*,***K132C)***

For labelling the protein RepA-WH1(A31V) with Alexa Fluor 488, a variant with only one cysteine was constructed. The gen *repA*-WH1(A31V) was amplified from a mutant variant in which Cys29 and Cys106 had been replaced by serines (Diederix et al., 2008), and a new C-terminal cysteine was added replacing Lys in position 132, using the following primer: 5’-GCTAAGCTTCAACAGAGCTGATAGCTGGTGAACTC. The *repA*-*WH1(A31V,K132C)* gene was interchanged with the insert in pET3d-*repA-WH1* (Fernández and Giraldo, 2018) upon digestion with SacII and BamHI.

### Protein Biochemistry

#### RepA-WH1(A31V) protein purification

The RepA-WH1(A31V) protein used in the fibrillation studies was purified as described in (Giraldo, 2007). RepA-WH1(A31V, K132C) was expressed in *E. coli* BL21(DE3) bearing pET-*WH1(A31V,K132C)*, plus pLysS to enable endogenous lysozyme-promoted lysis after freezing–thawing. Cells were grown in Terrific Broth medium supplemented with ampicillin (Ap) to 100 µg·ml^−1^ at 30 °C to an OD_600nm_ of 0.8. Then, IPTG was supplied to 0.5 mM and expression proceeded for 6 h at room temperature (RT). Cells were harvested and protocol was followed as described (Giraldo, 2007). The concentration of purified protein was determined by absorption at 280 nm (ε^280^ = 11,548 M^−1^·cm^−1^).

#### Fluorescent labelling of RepA-WH1

The single cysteine residue in RepA-WH1(A31V,K132C) was labelled with Alexa Fluor 488 maleimide (Invitrogen) using thiol chemistry. Briefly, the protein (~50 µM) was dialyzed against 20 mM of sodium phosphate buffer pH 7.0 and 100 mM Na_2_SO_4_. After dialysis, TCEP [tris(2-carboxyethyl) phosphine-hydrochloride] to 2 mM and urea to 2 M were added and incubated for 1 h on ice. TCEP maintains the cysteine residue in a reduced form, available for derivatization. The fluorophore was prepared by dissolving 250 µg in 10 µl H_2_O and added dropwise to the protein solution while gently agitated in a small vial. We used 5-fold molar excess of the dye to protein at 50 µM. From this point onwards, the sample was protected from light using aluminium foil. The conjugation reaction was allowed to proceed at RT for 4 h under mild agitation. After protein labelling, unconjugated Alexa Fluor 488 was separated in a 5 mL desalting column (GE Healthcare). Sample was divided into aliquots, flash frozen in N_2_(l), and stored at −70°C after adding glycerol to 10%. Each aliquot was thawed immediately before the experiment and used only once. The fraction of protein labelled was 20% (i.e., ~0.2 moles of dye per mole of protein).

#### Fibre assembly

Fibres were prepared with a modification of the protocol described in (Giraldo et al., 2007; Torreira et al., 2015). 25 µM of RepA-WH1(A31V) and 0.5 µM of Alexa 488-labelled RepA-WH1(A31V,K132C) were mixed in a final reaction volume of 100 µl of 0.1 M Na_2_SO_4_, 4 mM MgSO_4_, 40 mM HEPES pH 8 and 7% PEG-4000. 1 µl (diluted 1:100 from the stock) of purified intracellular RepA-WH1(A31V)-mCherry inclusions were added to nucleate amyloid assembly (Fernández-Tresguerres et al., 2010). Fibres were formed in 500 µl Eppendorf tubes at 4°C for 20 days. For cellular uptake experiments, fibres were centrifuged at 13,000 rpm for 30 min. After centrifugation, the supernatant was removed and fibres were washed with buffer (50 mM Na_2_SO_4_, 40 mM HEPES pH 8) and centrifuged again to be finally resupended in the same buffer. To get a more efficient internalization, immediately before incubation with cells, fibres were fragmented by 5 min sonication in a water bath (Ultrasonic cleaner USC100T, 60W), and further broken up by pipetting with a 100 μl Hamilton syringe for 2 min. Fibres were checked by means of electron microscopy (Fig. S2A), as indicated in (Giraldo, 2007; Fernández et al., 2016b), and confocal microscopy (Fig. S2B; see below). The size distribution of the fibres as added to the cell cultures was 257.6 ± 188.0 nm (n= 243).

#### Whole cell protein extraction

Either transiently transfected N2a or stable SH-SY5Y cells (see below) were rinsed in PBS, detached by scraping and harvested by centrifugation at 2,000 rpm, for 5 min at 4°C. Cell pellets were resuspended in 100 µl of RIPA buffer (Sigma), supplemented with protease and phosphatase inhibitors (Roche), incubated on ice for 30 min and then centrifuged at 14,000 rpm for 5 min at 4 °C. Pellets and supernatants were kept frozen at −80°C. Cell lysate supernatants were denatured with Laemmli buffer for 5 min at 95 °C and samples were loaded (30 µg total protein per lane) onto 12.5% SDS-denaturing polyacrylamide gels. Western blot analysis was performed after semi-dry transference of SDS-PAGE gels to PVDF membranes, using mouse anti-GFP (1:1,000; Abcam), or rabbit anti-mCherry (1:5,000; Abcam) antibodies, and anti-actin (1:1,000; Abcam) or anti-vinculin (1:1,000; Sigma) antibodies as loading controls. Then membranes were incubated with HRP-conjugated secondary anti-mouse/rabbit antibodies (1:10,000; Sigma). Blots were developed with ECL (Pierce).

### Cell Biology

#### Cell cultures

The murine neuroblastoma cell line Neuro 2a (N2a; DSMZ-Germany), and the human HeLa (DSMZ) and neuroblastoma SH-SY5Y (CIB-CSIC stock) cell lines were cultured in Dulbecco’s Modified Eagle Medium: nutrient Mixture F-12 (DMEM-F12) (Gibco) supplemented with 10% tetracycline-free fetal bovine serum (FBS) (Capricorn), 200 mM glutamine (Gibco) and 1% penicillin/streptomycin (Gibco). For the selection and maintenance of SH-SY5Y stable clones, geneticin (Gibco) and hygromycin B (Clontech) were added to the medium. Tet-ON expression was induced by doxycycline (Sigma). Cells culture was carried out at 37 °C in a humidified 5% CO_2_ incubator. Every 3-4 days, cells were grown to near confluence (80-90% density) and then detached using 0.05% trypsin-EDTA (Gibco) for subculturing.

#### N2a cells transfection

N2a cells were transiently transfected at ≈70% confluence in a 6-well plate by adding per well 10 μL Lipofectamine 2000 (Invitrogen) together with 3 μg of the respective pcDNA3.1 plasmid derivative. DNA-Lipofectamine complexes were prepared according to the manufacturer’s protocol in serum-free DMEM-F12 medium. Protein expression was followed by Western-blot analysis (Fig. S1B; see above) and confocal microscopy (see below).

#### Uptake of amyloid fibrils by N2a cells

To assay the uptake of the *in vitro* pre-assembled RepA-WH1(A31V)-Alexa 488 labelled fibres by the N2a cell line, cells were seeded in 35 mm plates and transiently transfected with pcDNA3.1-*repA-WH1(WT)-mCherry*, expressing a soluble variant of RepA-WH1 (Fernández-Tresguerres et al., 2010; Molina-García and Giraldo, 2014), or pcDNA3.1*-mCherry* as a control (see above; Fig. S1A). Immediately after transfection, N2a cells were exposed to RepA-WH1(A31V)-Alexa 488 fibres (1 μM in equivalent protein monomers) in the growth medium. Next day, cells were extensively rinsed with PBS and further cultured in fresh medium. Fluorescence was subsequently monitored by confocal microscopy 24, 48 and 72 h after the exposure to the fibres.

#### Generation of stable and Tet-ON regulated expression cell lines

To stably express RepA-WH1(WT)-TagGFP2 and TagGFP2 in the SH-SY5Y cell line, the Tet-ON 3G expression system (Clontech) was used. It is composed of pCMV-*Tet3G* (regulatory) and pTRE3G (response) plasmids. To generate SH-SY5Y cells expressing the trans-activator protein (rtTA) upon genomic integration of pCMV-*Tet3G*, 1 μg of this plasmid was transfected into 5×10^5^ SH-SY5Y cells with Lipofectamine LTX PLUS (1:1 ratio; Invitrogen) and selected for colonies with G418 (0.5 mg/ml). Individual clones, isolated using a cloning cylinder (Sigma), were then screened for luciferase activity (see below). Suitable THN-rtTA/SH-SY5Y clones (i.e., those with high luciferase induction and low basal expression) were subsequently co-transfected with a mixture (10:1) of pTRE3G-*WH1(WT)-TagGFP2* (or pTRE3G-*TagGFP2*) and a hygromycin linear selection marker. Double-stable SH-SY5Y transfectants were then selected by screening with hygromycin (0.4 mg/ml) and G418 (0.5 mg/ml) for 2 weeks, testing for the best expression of *repA-WH1(WT)-TagGFP2* and *TagGFP2* genes in the presence/absence of doxycycline (0.5 µg/ml) after 48 h by Western-blot (Fig. S3B; see above), flow cytometry (Fig. S3C) and confocal microscopy (Fig. S3D) (see below).

#### Luciferase assay

The luciferase assay kit (Promega) was used to test the expression and regulation performance of the selected transactivator-positive clones. Specifically, G418 resistant THN-rtTA/SH-SY5Y clones were seeded in a 6-well culture plate at a density of 5×10^5^ cells/well. One day after seeding, cells were transiently transfected with 5 µg of the pTRE3G-Luc control vector (Clontech) using Lipofectamine LTX (see above). Luciferase expression was induced by adding 0.5 µg/ml doxycycline for 48 h to complete DMEM-F12 medium. In parallel, to test clones for basal luciferase expression, controls with no doxycycline supplied were casted. Thereafter, cells were harvested in RIPA lysis buffer and bioluminescence was monitored at 562 nm using a TD-20/20 Turner Designs luminometer.

#### N2a/SH-SY5Y co-cultures

To explore cell-to-cell transmissibility of protein particles, SH-SY5Y stable cells (recipient cells), expressing either soluble WH1(WT)-TagGFP2 or TagGFP2, were seeded on glass coverslips coated with 0.1 mg/ml poly-L-lysine (Sigma) and casted in 24-well plates. Twenty-four h afterwards, N2a cells (donor cells) transiently transfected with pcDNA3.1-*WH1(A31V)-mCherry* or pcDNA3.1-*WH1(WT)-mCherry* were detached with trypsin and transferred to the plates bearing the SH-SY5Y receptor cells (2:1 ratio) and co-cultured for 5 days in DMEM-F12 completed media with 0.5 µg/ml doxycycline. Cells were then fixed with 4% p-formaldehyde (PFA) during 15 min and stained with DAPI (5 min, 1 μg/ml in PBS; Merck-Millipore).

#### Flow cytometry

Flow cytometry was used to monitor fluorescent protein expression in the stable clones for rtTA transactivator, either in the presence or absence of 0.5 µg/ml doxycycline, at 48 h after induction. Cells were detached by trypsinization from 35 mm culture dishes and centrifuged at 1,000 rpm for 5 min at 4°C. Then, pellets were resuspended in 2 ml of cold PBS buffer. GFP (Excitation/Ex 488 nm; Emission/Em 525 nm) fluorescence was measured in a Coulter Epics XL (Beckman-Coulter) flow cytometer, analysing 10,000 cells per sample. Subsequent data analysis was performed using the Flowlogic software (v7.2.1).

#### Cell proliferation (MTT) assay

To assess cell proliferation, N2a cells transfected with pcDNA3.1-*WH1(Δ/WT/A31V/ΔN37)-mCherry*, as described above, were seeded 24 h after transfection into 96-well culture plates at a density of 6×10^4^ cells/well. The next day, MTT (thiazolyl blue tetrazolium bromide, Sigma; 0.5 mg/ml in PBS) was supplied and, after 4 h of incubation at 37°C, MTT was removed and replaced with 200 µl of DMSO. Absorbance was measured at 570 nm in a Varioskan Flash plate reader (Thermo Fisher Scientific). Absorbance values were normalized to those obtained in DMSO.

#### Confocal microscopy

Cells were directly observed at RT by confocal laser microscopy using a Leica TCS-SP5 microscope with a 63x (NA = 1.4–0.60/Oil HCX PL APO) immersion objective. Laser lines used for excitation were 514 nm (mCherry), 488 nm (Alexa 488 and Tag GFP2) and 405 nm (DAPI and ThS). Images were acquired every 1 μm (Z-sections). Cells were fixed with 4% PFA in PBS for 15 min, rinsed three times in PBS and nuclei staining, when required, was performed with 1 µg/ml DAPI solution (Merck-Millipore) in PBS for 10 min, then rinsed three times with PBS. Coverslips were mounted with Fluoromount-G medium (SouthernBiotech) on microscopic slides (Linealab). For amyloid characterization of intracellular particles, transiently transfected N2a cells were grown on coverslips coated with poly-L-lysine for 48 h. Cells were fixed with 4% PFA in PBS for 20 min and rinsed as above. Then, cells were permeabilized with cold 50% methanol (Emsure) in Milli-Q H_2_O for 5 min, incubated with 0.05% thioflavin-S (ThS, Sigma) in 12.5% ethanol for 30 min and rinsed three times in 50% ethanol in destilled H_2_O. Cells were then hydrated for 5 min in PBS and mounted.

#### Statistical analysis

All experiments were carried out in triplicate. GraphPath PRISM (v.6) software was used to estimate the media and standard deviation of data points, and for the analysis of their statistical significance, either through the Student’s *t*-test or one-way ANOVA.

### Proteomics

#### LC-MS analysis

The following cell types were studied: i) N2a cells transfected with pcDNA3.1-*WH1(WT)-mCherry* or pcDNA3.1-*mCherry*, and incubated with *in vitro*-assembled WH1(A31V) fibres for 48 h; ii) THN-rtTA/SH-SY5Y cells having integrated pTRE3G-*WH1(WT)-TagGFP2* or pTRE3G*-TagGFP2*, both after induction with Dox (0.5 µg/ml) for 48 h; and iii) naïve N2a and THN-rtTA/SH-SY5Y control cells co-cultivated as in (i) and (ii). Whole cell extracts (see above) were partially resolved by SDS-PAGE gel (10% polyacrylamide), so that the whole proteome became concentrated in the stacking/resolving gel interface. The unresolved protein bands were visualized by Colloidal Blue Staining (Invitrogen), excised into pieces (1 x 1 mm), and placed in 0.5 ml tubes and subjected to manual tryptic digestion (Cristobo et al., 2001). The eluted tryptic peptides were dried by speed-vacuum centrifugation and then desalted through ZipTip C18 Pipette tips (Millipore/Sigma-Aldrich). Samples were reconstituted in 10 µL of 0.1% formic acid before their analysis by nLC–MS/MS. All peptide separations were carried out on an Easy-nLC 1000 nanosystem (Thermo Fisher Scientific). For each analysis, the sample was loaded into an Acclaim PepMap 100 precolumn (Thermo Fisher Scientific) and eluted through a RSLC PepMap C18 column (50 cm long, 75 µm inner diameter and 2 µm particle size; Thermo Scientific). The mobile phase was 0.1% formic acid in water (solvent A) and 0.1% formic acid in acetonitrile (solvent B). The gradient profile was set, at a flow rate of 300 nL/min, as follows: 5–35% solvent B for 100 min, 35%-45% solvent B for 20 min, 45%-100% solvent B for 5 min, and 100% solvent B for 15 min. Four microliters of each sample were injected. MS analysis was performed using a Q-Exactive mass spectrometer (Thermo Scientific). Ionization was performed at 2,000 V of liquid junction voltage and a capillary temperature of 270 °C. The full scan method comprised a m/z 300–1800 mass selection, an Orbitrap resolution of 70,000 (at m/z 200), a target automatic gain control (AGC) value of 3×10^6^, and 100 ms of maximum injection time. After the survey scan, the 15 most intense precursor ions were selected for MS/MS fragmentation. Fragmentation was performed with a normalized collision energy of 27 and MS/MS scans were acquired with a starting mass of m/z 200, 2×10^5^ AGC target, a resolution of 17,500 (at m/z 200), 8×10^3^ of intensity threshold, 2.0 m/z units as isolation window and 100 ms maximum IT. Charge state screening was enabled to reject unassigned, singly charged and ≥ 7 protonated ions. A dynamic exclusion time of 30 s was used to discriminate against previously selected ions.

#### MS data analysis

MS data were analyzed with Proteome Discoverer (version 1.4.1.14) (Thermo) using standardized workflows. Mass spectra *.raw files were searched either against the *Mus musculus* SwissProt 2016 database (16,838 protein entries; for the N2a recipient cells), or the *Homo sapiens* SwissProt 2016 database (20,131 protein entries; for the SH-SY5Y recipient cells), using the Mascot search engine (v.2.6, Matrix Science). In the latter case, limiting the survey to the human proteome excluded proteins potentially coming from the donor murine cells. Precursor and fragment mass tolerance were set to 10 ppm and 0.02 Da, respectively, allowing 2 missed cleavages, carbamido methylation of cysteines as a fixed modification, and methionine oxidation and acetylation N-terminal as a variable modification. Identified peptides were filtered using Percolator algorithm with a q-value threshold of 0.01 (High Confidence Filter settings, FDR < 1%) (Käll et al., 2007). Proteins contributing peptides detected at least two times (PSM >1) or having more than one peptide identified were selected for further analysis. Proteins identified in naïve N2a murine control cells when co-cultured with fibres were subtracted from those found in N2a cells expressing WH1(WT)-mCherry or mCherry and also incubated with fibres. Proteins found in naïve human THN-rtTA/SH-SY5Y control cells co-cultured with murine cells donating WH1(A31V)-mCherry were subtracted from those in THN-rtTA/SH-SY5Ycells expressing WH1(WT)-TagGFP2 or TagGFP2 and also co-cultured with the murine donor cells. A Boolean algebra analysis (Venny, v.2.1; https://bioinfogp.cnb.csic.es/tools/venny/) was undertaken to identify those host proteins found differentially under each experimental condition. The subset of proteins present in the WH1(WT)-mCherry/TagGFP2 datasets solely upon incubation with the WH1(A31V) fibres/donor cells were then classified according to functional gene ontology (GO) (GeneCards; Stelzer et al., 2010) and cluster analysis (STRING v.11; Szklarczyk et al., 2019).

## Acknowledgements

The authors thank their groups for much support and encouragement. We are grateful to Ana Serrano for help with plasmid construction and to the CIB-CSIC staff members Carmen Doñoro (cell culture facility) and Maite Seisdedos and Gema Elvira (confocal microscopy service). The help of Pedro Lastres (CIB-CSIC) and Berta Raposo (CBM-CSIC/UAM) with flow cytometry is also acknowledged. We are indebted to Patricia Boya and her group and to Patricio Aller for helpful advice and discussions. This work has been financed with grants from the Spanish AEI (BIO2012-30852 and BIO2015-68730-R) to RG. ARG was a recipient of an AEI short-term fellowship (EEBB-I-17-12294) to work at the laboratory of IMV.

## Author contributions

ARG carried out most experimental work and analysed data; CF prepared the fluorescence-labelled fibres and helped with their transfection and data analysis; MMdA made the plasmid constructs and earlier work with the SH-SY5Y cells; VdlR performed the proteomic analyses; IMV hosted ARG and provided expertise on the cell biology of cytoplasmic propagation of prion-like proteins; RG conceived the project, analysed data and wrote the manuscript with contributions from all the authors.

## Conflict of interest

The authors declare that they have no conflict of interest.

## Supplementary Files

**Fig. S1.**
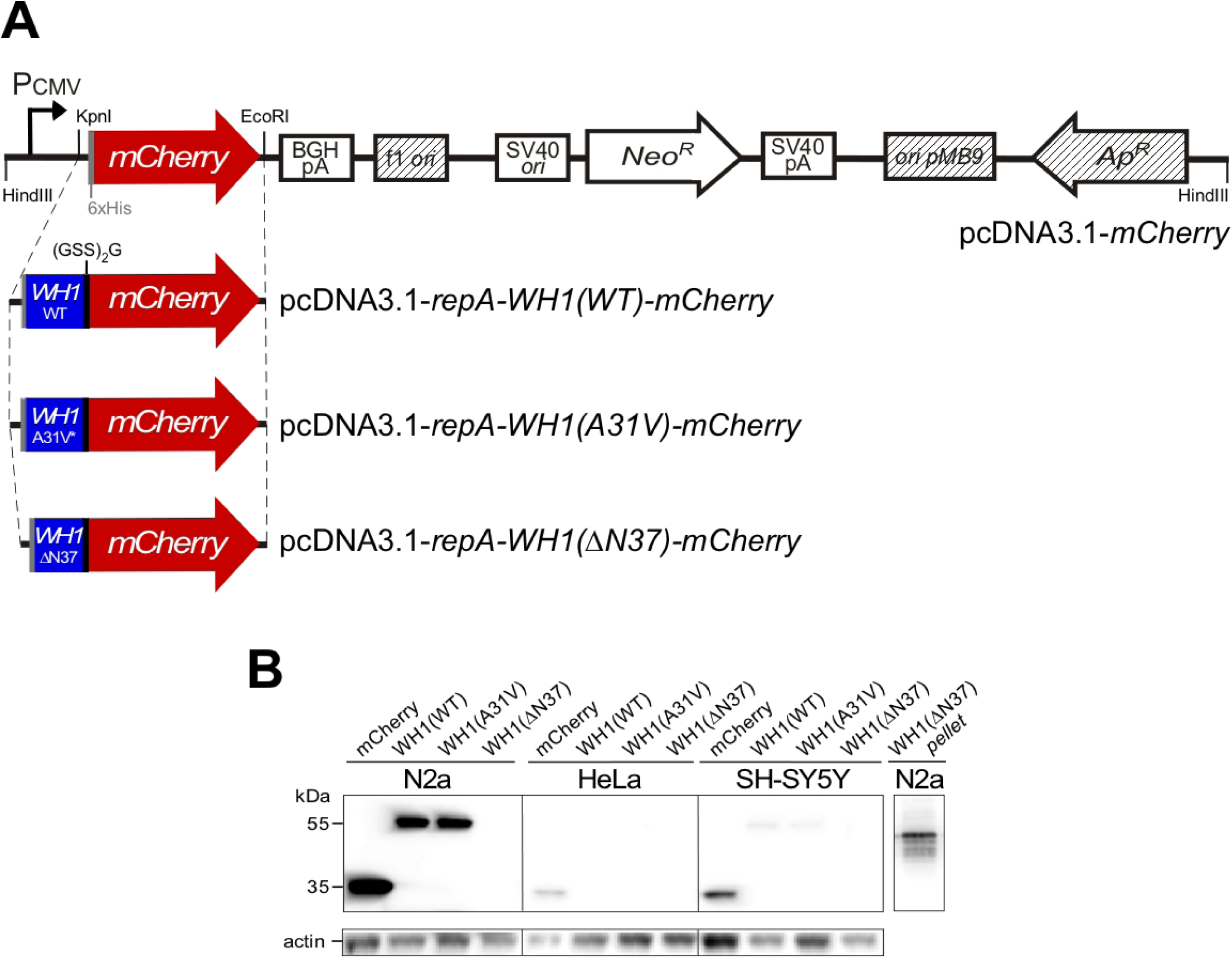
Transient expression in mammalian cell lines of distinct variants of the bacterial prion-like protein RepA-WH1 fused to the red fluorescent protein mCherry. (**A**) Schematic linear representation of the pcDNA3.1 plasmid constructs used to transiently express either the distinct WH1-mCherry fusions or the mCherry control in mammalian cell lines. (**B**)Assessing the expression of the WH1-mCherry derived constructions (WT/A31V/ΔN37; ≈55 kDa), or the mCherry control (35 kDa), in (A) by Western-blotting of soluble lysates 48 h after transient transfection of N2a, SH-SY5Y and HeLa cell lines. Protein constructions were detected using an anti-mCherry antibody. Actin (42 kDa) as a loading control was visualized with an anti-actin antibody. The insoluble fraction (pellet) of cells expressing WH1(ΔN37) in the N2a cells is also shown, compatible with its tendency to form large foci in bacteria (Gasset-Rosa et al., 2014; Molina-García et al., 2014).

**Fig. S2.**
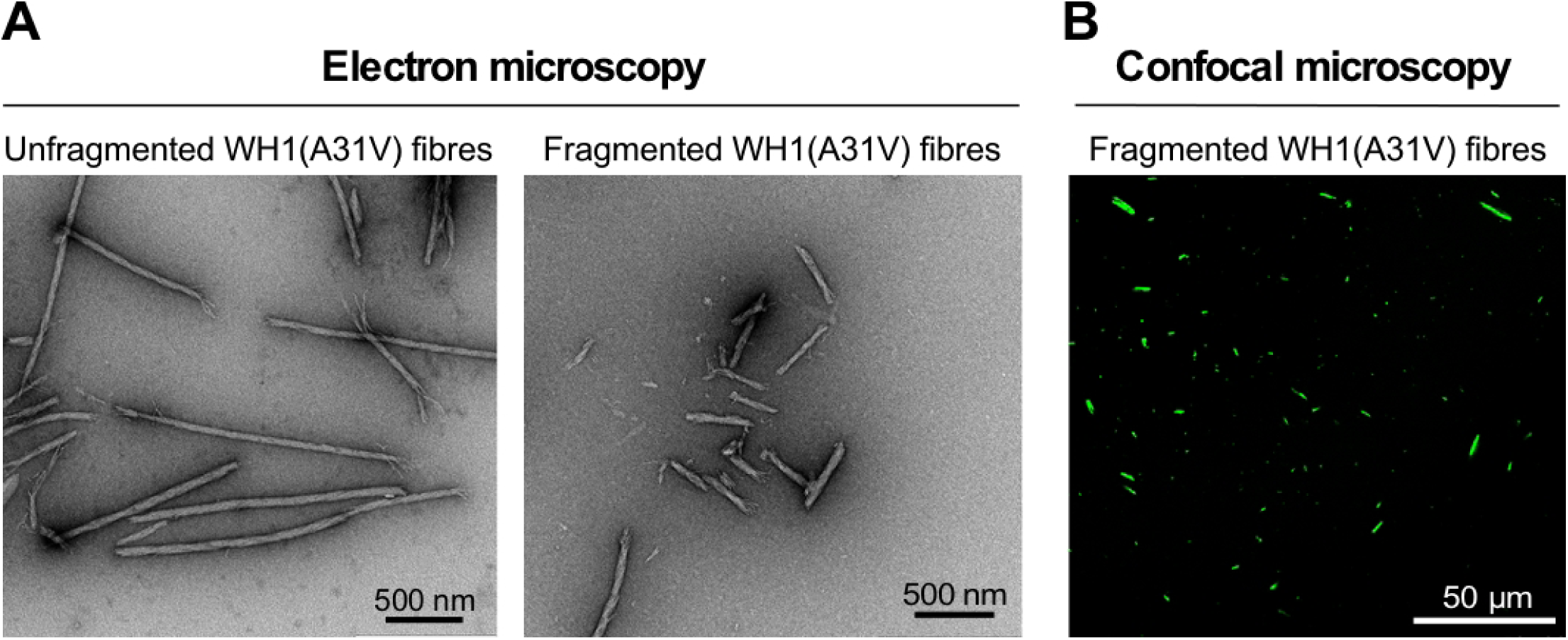
*In vitro*-assembled and Alexa 488-labelled WH1(A31V) fibres used for co-cultivation with the murine N2a cells. (**A**) Negative staining TEM micrographs of the fibres before (*left*) and after (*right*) being disrupted by means of sonication, previously to their addition to the transfected N2a cells. (**B**) Confocal section of the fibre samples. Green fluorescence from the tracer Alexa 488-labelled WH1(A31V) molecules incorporated into the fibres is evident.

**Fig. S3.**
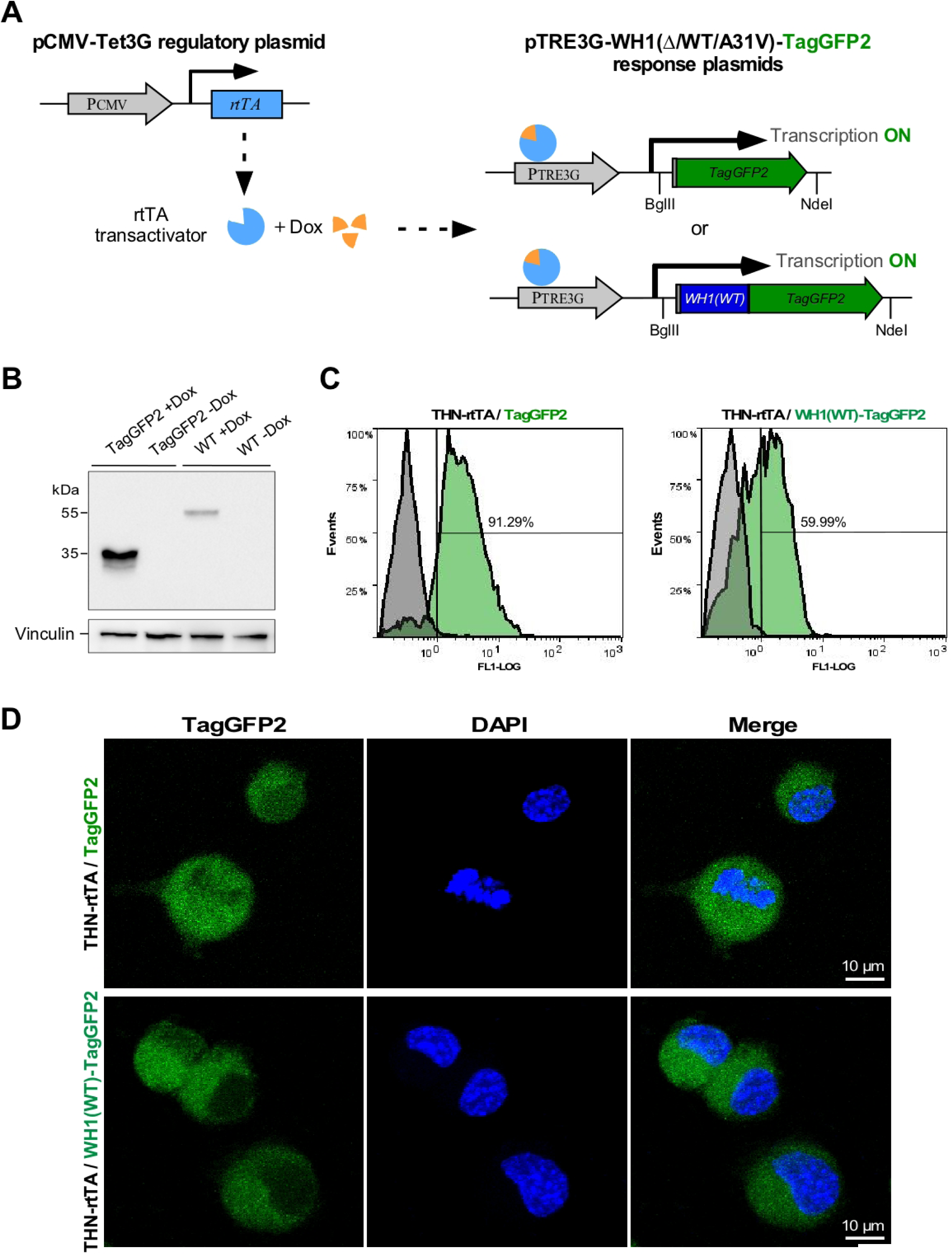
Experimental setting used for the stable expression of proteins in the human neuroblastoma cell line SH-SY5Y. (**A**) Tet-ON, doxycline (Dox)-dependent expression cascade from the integrated pCMV-Tet3G transactivator (rtTA; *left*) and the pTRE3G response plasmids (*right*). (**B**) Western-blots of protein extracts from THN-rtTA/SH-SY5Y cells expressing WH1(WT)-TagGFP2 or TagGFP2 upon induction for 48 h with 0.5 µg/ml Dox. Protein fusions were probed with an anti-GFP antibody, whereas detection of vinculin (≈123 kDa) was used as a loading control. (**C**) Flow cytometry histograms (FL1 channel) highlighting the population of live THN-rtTA/SH-SY5Y cells expressing (green) or not (grey) the engineered fluorescent proteins after 48 h of culture in the presence of Dox (see B). Mean fluorescence intensity (MFI) TagGFP2: 3.44; MFI WH1(WT)-TagGFP2: 1.61. (**D**) Confocal maximum-intensity projections images of THN-rtTA/SH-SY5Y clones stably expressing the rtTA transactivator and TagGFP2, or WH1(WT)-TagGFP2 (green), 48 h after Dox addition. Nuclei were visualized using DAPI (blue). Both proteins are expressed soluble (diffuse fluorescence) in the cytoplasm.

**Fig. S4.**
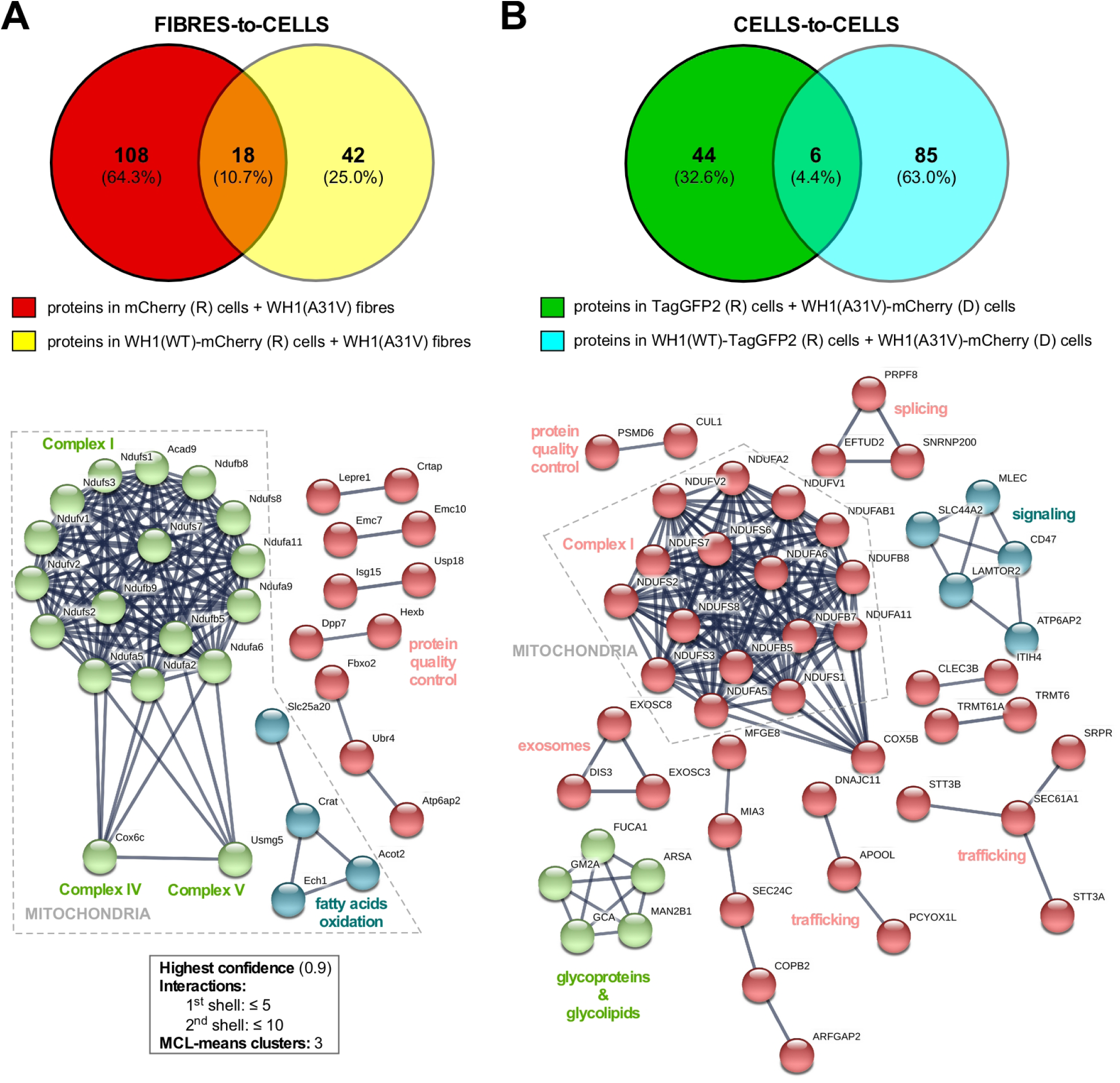
Systems analysis of the response of murine N2a (**A**) and human THN-rtTA/SH-SY5Y (**B**) cells, expressing soluble WH1(WT) or its fluorescent tags, to co-cultivation with the *in vitro*-assembled and the cell-released WH1(A31V) amyloids, respectively. Venn diagrams (*top*) indicate the number of proteins exclusively identified in the WH1(WT)-mCherry/TagGFP2 or in the control mCherry/TagGFP2 cells (Figure 6), as well as in both of them, over those found in the naïve N2a or THN-rtTA/SH-SY5Y backgrounds when challenged with the same fibres or donor cells, which were subtracted. STRING v.11 (Szklarczyk et al., 2019) functional clustering analysis (*bottom*; inset: program settings) was performed, using all available prediction sources, for the proteins identified as differentially expressed under both experimental settings. After removing the unconnected nodes, the remaining clusters, similar under both amyloid transmission scenarios, point to contributions of the mitochondrial membrane respiratory complexes, protein quality control and signalling/trafficking to the RepA-WH1 ‘generic’ model of amyloidosis in mammalian cells.

**Supplemental Dataset S1.** ESI-MS/MS analysis of the proteome response of mouse N2a cells to WH1(A31V) fibril uptake-promoted amyloidogenesis.

**Supplemental Dataset S2.** ESI-MS/MS analysis of the proteome response of human THN-rtTA/SH-SY5Y cells on transmission of WH1(A31V)-mCherry from murine N2a cells.

**Table S1.** Subset of proteins in Datasets S1 and S2 matching proteins/genes differentially located in published system analyses of aggregation-triggered neurodegeneration.

## References

Aguzzi A, Falsig J. 2012. Prion propagation, toxicity and degradation. Nature Neuroscience 15:936–939. DOI: 10.1038/nn.3120.

Aguzzi A, Lakkaraju AKK. 2016. Cell Biology of Prions and Prionoids: A Status Report. Trends in Cell Biology 26:40–51. DOI: 10.1016/j.tcb.2015.08.007.

Chiti F, Dobson CM. 2017. Protein Misfolding, Amyloid Formation, and Human Disease: A Summary of Progress Over the Last Decade. Annual Review of Biochemistry 86:27–68. DOI: 10.1146/annurev-biochem-061516-045115.

Clavaguera F, Bolmont T, Crowther RA, Abramowski D, Frank S, Probst A, Fraser G, Stalder AK, Beibel M, Staufenbiel M, Jucker M, Goedert M, Tolnay M. 2009. Transmission and spreading of tauopathy in transgenic mouse brain. Nature Cell Biology 11:909–913. DOI: 10.1038/ncb1901.

Cristobo I, Larriba MJ, de los Ríos V, García F, Muñoz A, Casal JI. 2011. Proteomic analysis of 1α,25-dihydroxyvitamin D3 action on human colon cancer cells reveals a link to splicing regulation. J Proteomics 75:384–397. DOI: 10.1016/j.jprot.2011.08.003.

de Villiers EM, Gunst K, Chakraborty D, Ernst C, Bund T, Zur Hausen H. 2019. A specific class of infectious agents isolated from bovine serum and dairy products and peritumoral colon cancer tissue. Emerging Microbes & Infections 8:1205–1218. DOI: 10.1080/22221751.2019.1651620.

Diederix RE, Dávila C, Giraldo R, Lillo MP. 2008. Fluorescence studies of the replication initiator protein RepA in complex with operator and iteron sequences and free in solution. FEBS Journal 275:5393–5407. DOI: 10.1111/j.1742-4658.2008.06669.x.

Duernberger Y, Liu S, Riemschoss K, Paulsen L, Bester R, Kuhn PH, Schölling M, Lichtenthaler SF, Vorberg I. 2018. Prion replication in the mammalian cytosol: Functional regions within a prion domain driving induction, propagation, and inheritance. Molecular and Cellular Biology 38:e00111–18. DOI: 10.1128/MCB.00111-18.

Eisele YS, Obermüller U, Heilbronner G, Baumann F, Kaeser SA, Wolburg H, Walker LC, Staufenbiel M, Heikenwalder M, Jucker M. 2010. Peripherally applied Abeta-containing inoculates induce cerebral beta-amyloidosis. Science 330:980–982. DOI: 10.1126/science.1194516.

Eisenberg DS, Sawaya MR. 2017. Structural studies of amyloid proteins at the molecular level. Annual Review of Biochemistry 86:69–95. DOI: 0.1146/annurev-biochem-061516-045104.

Fernández C, Giraldo R. 2018. Modulation of the aggregation of the prion-like protein RepA-WH1 by chaperones in a cell-free expression system and in cytomimetic lipid vesicles. ACS Synthetic Biology 7:2087–2093. DOI: 10.1021/acssynbio.8b00283.

Fernández C, González-Rubio G, Langer J, Tardajos G, Liz-Marzán LM, Giraldo R, Guerrero-Martínez A. 2016a. Nucleation of amyloid oligomers by RepA-WH1 prionoid-functionalized gold nanorods. Angewandte Chemie International Edition 55:11237–11241. DOI: 10.1002/anie.201604970.

Fernández C, Núñez-Ramírez R, Jiménez M, Rivas G, Giraldo R. 2016b. RepA-WH1, the agent of an amyloid proteinopathy in bacteria, builds oligomeric pores through lipid vesicles. Scientific Reports 6:23144. DOI: 10.1038/srep23144.

Fernández-Tresguerres ME, Moreno-Díaz de la Espina S, Gasset-Rosa F, Giraldo R. 2010. A DNA-promoted amyloid proteinopathy in *Escherichia coli*. Molecular Microbiology 77:1456–1469. DOI: 10.1111/j.1365-2958.2010.07299.x.

Frederick KK, Debelouchina GT, Kayatekin C, Dorminy T, Jacavone AC, Griffin RG, Lindquist S. 2014. Distinct prion strains are defined by amyloid core structure and chaperone binding site dynamics. Chemistry & Biology 21:295–305. DOI: 10.1016/j.chembiol.2013.12.013.

Friedland RP, Chapman MR. 2017. The role of microbial amyloid in neurodegeneration. PLoS Pathogens 13:e1006654. DOI 10.1371/journal.ppat.1006654.

Frost B, Jacks RL, Diamond MI. 2009. Propagation of tau misfolding from the outside to the inside of a cell. Journal of Biological Chemistry 284:12845–12852. DOI: 10.1074/jbc.M808759200.

Garrity SJ, Sivanathan V, Dong J, Lindquist S, Hochschild A. 2010. Conversion of a yeast prion protein to an infectious form in bacteria. Proceedings of the National Academy of Sciences of the United States of America 107:10596–10601. DOI: 10.1073/pnas.0913280107.

Gasset-Rosa F, Coquel AS, Moreno-del Álamo M, Chen P, Song X, Serrano AM, Moreno-Díaz de la Espina S, Lindner AB, Giraldo R. 2014. Direct assessment in bacteria of prionoid propagation and phenotype selection by Hsp70 chaperone. Molecular Microbiology 91:1070–1087. DOI: 10.1111/mmi.12518.

Gasset-Rosa F, Giraldo R. 2015. Engineered bacterial hydrophobic oligopeptide repeats in a synthetic yeast prion, [*REP-PSI*^+^]. Frontiers in Microbiology 6:311. DOI: 10.3389/fmicb.2015.00311.

Giraldo R, Fernández-Tornero C, Evans PR, Díaz-Orejas R, Romero A. 2003. A conformational switch between transcriptional repression and replication initiation in the RepA dimerization domain. Nature Structural Biology 10:565–571. DOI: 10.1038/nsb937.

Giraldo R. 2007. Defined DNA sequences promote the assembly of a bacterial protein into distinct amyloid nanostructures. Proceedings of the National Academy of Sciences of the United States of America 104:17388–17393. DOI: 10.1073/pnas.0702006104.

Giraldo R. 2019. Optogenetic navigation of routes leading to protein amyloidogenesis in bacteria. Journal of Molecular Biology 431:1186–1202. DOI: 10.1016/j.jmb.2019.01.037.

Goedert M. 2015. Alzheimer’s and Parkinson’s diseases: The prion concept in relation to assembled Aβ, tau, and α-synuclein. Science 349:1255555. DOI: 10.1126/science.1255555.

Goedert M, Eisenberg DS, Crowther RA. 2017. Propagation of Tau Aggregates and Neurodegeneration. Annual Review of Neuroscience 40:189–210. DOI: 10.1146/annurev-neuro-072116-031153.

Haelterman NA, Yoon WH, Sandoval H, Jaiswal M, Shulman JM, Bellen HJ. 2014. A mitocentric view of Parkinson’s disease. Annual Review Neuroscience 37:137–159. doi: 10.1146/annurev-neuro-071013-014317.

Hofmann JP, Denner P, Nussbaum-Krammer C, Kuhn PH, Suhre MH, Scheibel T, Lichtenthaler SF, Schätzl HM, Bano D, Vorberg IM. 2013. Cell-to-cell propagation of infectious cytosolic protein aggregates. Proceedings of the National Academy of Sciences of the United States of America 110:5951–5956. DOI: 10.1073/pnas.1217321110.

Hosp F, Vossfeldt H, Heinig M, Vasiljevic D, Arumughan A, Wyler E, GERAD1 Consortium, Landthaler M, Hubner N, Wanker EE, Lannfelt L, Ingelsson M, Lalowski M, Voigt A, Selbach M. 2015. Quantitative interaction proteomics of neurodegenerative disease proteins. Cell Reports 11:1134–1146. DOI: 10.1016/j.celrep.2015.04.030.

Hwang D, Lee IY, Yoo H, Gehlenborg N, Cho JH, Petritis B, Baxter D, Pitstick R, Young R, Spicer D, Price ND, Hohmann JG, Dearmond SJ, Carlson GA, Hood LE. 2009. A systems approach to prion disease. Molecular Systems Biology 5:252. DOI: 10.1038/msb.2009.10.

Jucker M, Walker LC. 2018. Propagation and spread of pathogenic protein assemblies in neurodegenerative diseases. Nature Neuroscience 21:1341–1349. DOI: 10.1038/s41593-018-0238-6.

Käll L, Canterbury JD, Weston J, Noble WS, MacCoss MJ. 2007. Semi-supervised learning for peptide identification from shotgun proteomics datasets. Nature Methods 4:923–925. DOI: 10.1038/nmeth1113.

Kfoury N, Holmes BB, Jiang H, Holtzman DM, Diamond MI. 2012. Trans-cellular propagation of Tau aggregation by fibrillar species. Journal of Biological Chemistry 287:19440–19451. DOI: 10.1074/jbc.M112.346072.

Kilic T, Popov AN, Burk-Körner A, Koromyslova A, Zur Hausen H, Bund T, Hansman GS. 2019. Structural analysis of a replication protein encoded by a plasmid isolated from a multiple sclerosis patient. Acta Crystallographica D 75:498–504. DOI: 10.1107/S2059798319003991.

Kim YE, Hosp F, Frottin F, Ge H, Mann M, Hayer-Hartl M, Hartl FU. 2016. Soluble Oligomers of PolyQ-Expanded Huntingtin Target a Multiplicity of Key Cellular Factors. Molecular Cell 63:951–964. DOI: 10.1016/j.molcel.2016.07.022.

Kordower JH, Chu Y, Hauser RA, Freeman TB, Olanow CW. 2008. Lewy body-like pathology in long-term embryonic nigral transplants in Parkinson’s disease. Nature Medicine 14:504–506. DOI: 10.1038/nm1747.

Krammer C, Kryndushkin D, Suhre MH, Kremmer E, Hofmann A, Pfeifer A, Scheibel T, Wickner RB, Schätzl HM, Vorberg I. 2009. The yeast Sup35NM domain propagates as a prion in mammalian cells. Proceedings of the National Academy of Sciences of the United States of America 106:462–467. DOI: 10.1073/pnas.0811571106.

Liu S, Hossinger A, Hofmann JP, Denner P, Vorberg IM. 2016. Horizontal transmission of cytosolic Sup35 prions by extracellular vesicles. 2016. mBio 7:e00915–16. DOI: 10.1128/mBio.00915-16.

Lu JX, Qiang W, Yau WM, Schwieters CD, Meredith SC, Tycko R. 2013. Molecular structure of β-amyloid fibrils in Alzheimer’s disease brain tissue. Cell 154:1257–1268. DOI: 10.1016/j.cell.2013.08.035.

Luk KC, Kehm VM, Zhang B, O’Brien P, Trojanowski JQ, Lee VM. 2012. Intracerebral inoculation of pathological α-synuclein initiates a rapidly progressive neurodegenerative α-synucleinopathy in mice. Journal of Experimental Medicine 209:975–986. DOI: 10.1084/jem.20112457.

Luk KC, Song C, O’Brien P, Stieber A, Branch JR, Brunden KR, Trojanowski JQ, Lee VM. 2009. Exogenous alpha-synuclein fibrils seed the formation of Lewy body-like intracellular inclusions in cultured cells. Proceedings of the National Academy of Sciences of the United States of America 106:20051–20056. DOI: 10.1073/pnas.0908005106.

Markmiller S, Soltanieh S, Server KL, Mak R, Jin W, Fang MY, Luo EC, Krach F, Yang D, Sen A, Fulzele A, Wozniak JM, Gonzalez DJ, Kankel MW, Gao FB, Bennett EJ, Lécuyer E, Yeo GW. 2018. Context-Dependent and Disease-Specific Diversity in Protein Interactions within Stress Granules. Cell 172:590–604. DOI: 10.1016/j.cell.2017.12.032.

Mathys H, Davila-Velderrain J, Peng Z, Gao F, Mohammadi S, Young JZ, Menon M, He L, Abdurrob F, Jiang X, Martorell AJ, Ransohoff RM, Hafler BP, Bennett DA, Kellis M, Tsai LH. 2019. Single-cell transcriptomic analysis of Alzheimer’s disease. Nature 570:332–337. DOI: 10.1038/s41586-019-1195-2.

Molina-García L, Gasset-Rosa F, Moreno-del Álamo M, Fernández-Tresguerres ME, Moreno-Díaz de la Espina S, Lurz R, Giraldo R. 2016. Functional amyloids as inhibitors of plasmid DNA replication. Scientific Reports 6:25425. DOI: 10.1038/srep25425.

Molina-García L, Giraldo R. 2014. Aggregation interplay between variants of the RepA-WH1 prionoid in *Escherichia coli*. Journal of Bacteriology 196:2536–2542. DOI: 10.1128/JB.01527-14.

Molina-García L, Moreno-del Álamo M, Botias P, Martín-Moldes Z, Fernández M, Sánchez-Gorostiaga A, Alonso-del Valle A, Nogales J, García-Cantalejo J, Giraldo R. 2017. Outlining core pathways of amyloid toxicity in bacteria with the RepA-WH1 prionoid. Frontiers in Microbiology 8:539. DOI: 10.3389/fmicb.2017.00539.

Moreno-del Álamo M, Moreno-Díaz de la Espina S, Fernández-Tresguerres ME, Giraldo R. 2015. Pre-amyloid oligomers of the proteotoxic RepA-WH1 prionoid assemble at the bacterial nucleoid. Scientific Reports 5:14669. DOI: 10.1038/srep14669.

Norambuena A, Wallrabe H, Cao R, Wang DB, Silva A, Svindrych Z, Periasamy A, Hu S, Tanzi RE, Kim DY, Bloom GS. 2018. A novel lysosome-to-mitochondria signaling pathway disrupted by amyloid-β oligomers. EMBO Journal 37:e100241. DOI: 10.15252/embj.2018100241.

Olzscha H, Schermann SM, Woerner AC, Pinkert S, Hecht MH, Tartaglia GG, Vendruscolo M, Hayer-Hartl M, Hartl FU, Vabulas RM. 2011. Amyloid-like aggregates sequester numerous metastable proteins with essential cellular functions. Cell 144:67–78. DOI: 10.1016/j.cell.2010.11.050.

Perov S, Lidor O, Salinas N, Golan N, Tayeb-Fligelman E, Deshmukh M, Willbold D, Landau M. 2019. Structural Insights into Curli CsgA Cross-β Fibril Architecture Inspire Repurposing of Anti-amyloid Compounds as Anti-biofilm Agents. PLoS Pathogens 15:e1007978. DOI: 10.1371/journal.ppat.1007978.

Polymenidou M, Cleveland DW. 2012. Prion-like spread of protein aggregates in neurodegeneration. Journal of Experimental Medicine 209:889–893. DOI: 10.1084/jem.20120741.

Prusiner SB. 1998. Prions. Proceedings of the National Academy of Sciences of the United States of America 95:13363–13383.

Prusiner SB. 2013. Biology and genetics of prions causing neurodegeneration. Annual Review of Genetics 47:601–623. DOI: 10.1146/annurev-genet-110711-155524.

Prusiner SB, Woerman AL, Mordes DA, Watts JC, Rampersaud R, Berry DB, Patel S, Oehler A, Lowe JK, Kravitz SN, Geschwind DH, Glidden DV, Halliday GM, Middleton LT, Gentleman SM, Grinberg LT, Giles K. 2015. Evidence for α-synuclein prions causing multiple system atrophy in humans with parkinsonism. Proceedings of the National Academy of Sciences of the United States of America 112:E5308–E5317. DOI: 10.1073/pnas.1514475112.

Qiang W, Yau WM, Lu JX, Collinge J, Tycko R. 2017. Structural variation in amyloid-β fibrils from Alzheimer’s disease clinical subtypes. Nature 541:217–221. DOI: 10.1038/nature20814.

Riemschoss K, Arndt V, Bolognesi B, von Eisenhart-Rothe P, Liu S, Buravlova O, Duernberger Y, Paulsen L, Hornberger A, Hossinger A, Lorenzo-Gotor N, Hogl S, Müller SA, Tartaglia G, Lichtenthaler SF, Vorberg IM. 2019. Fibril-induced glutamine-/asparagine-rich prions recruit stress granule proteins in mammalian cells. Life Science Alliance 2:e201800280. DOI: 10.26508/lsa.201800280.

Seyfried NT, Dammer EB, Swarup V, Nandakumar D, Duong DM, Yin L, Deng Q, Nguyen T, Hales CM, Wingo T, Glass J, Gearing M, Thambisetty M, Troncoso JC, Geschwind DH, Lah JJ, Levey AI. 2017. A Multi-network Approach Identifies Protein-Specific Co-expression in Asymptomatic and Symptomatic Alzheimer’s Disease. Cell Systems 4:60–72. DOI: 10.1016/j.cels.2016.11.006.

Scheckel C, Aguzzi A. 2018. Prions, prionoids and protein misfolding disorders. Nature Reviews in Genetics 19:405–418. DOI: 10.1038/s41576-018-0011-4.

Sonati T, Reimann RR, Falsig J, Baral PK, O’Connor T, Hornemann S, Yaganoglu S, Li B, Herrmann US, Wieland B, Swayampakula M, Rahman MH, Das D, Kav N, Riek R, Liberski PP, James MN, Aguzzi A. 2013. The toxicity of antiprion antibodies is mediated by the flexible tail of the prion protein. Nature 501:102–106. DOI: 10.1038/nature12402.

Soto C. 2012. Transmissible proteins: expanding the prion heresy. Cell 149:968–977. DOI: 10.1016/j.cell.2012.05.007.

Stöhr J, Watts JC, Mensinger ZL, Oehler A, Grillo SK, DeArmond SJ, Prusiner SB, Giles K. 2012. Purified and synthetic Alzheimer’s amyloid beta (Aβ) prions. Proceedings of the National Academy of Sciences of the United States of America 109:11025–11030. DOI: 10.1073/pnas.1206555109.

Stelzer G, Rosen R, Plaschkes I, Zimmerman S, Twik M, Fishilevich S, Iny Stein T, Nudel R, Lieder I, Mazor Y, Kaplan S, Dahary D, Warshawsky D, Guan-Golan Y, Kohn A, Rappaport N, Safran M, Lancet D. 2020. The GeneCards Suite: From Gene Data Mining to Disease Genome Sequence Analysis. Current Protocols in Bioinformatics 54:1.30.1–1.30.33. DOI: 10.1002/cpbi.5.

Szklarczyk D, Gable AL, Lyon D, Junge A, Wyder S, Huerta-Cepas J, Simonovic M, Doncheva NT, Morris JH, Bork P, Jensen LJ, Mering CV. 2019. STRING v11: protein-protein association networks with increased coverage, supporting functional discovery in genome-wide experimental datasets. Nucleic Acids Research 47:D607–D613. DOI: 10.1093/nar/gky1131.

Torreira E, Moreno-del Álamo M, Fuentes-Perez ME, Fernández C, Martín-Benito J, Moreno-Herrero F, Giraldo R, Llorca O. 2015. Amyloidogenesis of bacterial prionoid RepA-WH1 recapitulates dimer to monomer transitions of RepA in DNA replication initiation. Structure 23:183–189. DOI: 10.1016/j.str.2014.11.007.

Wang P, Deng J, Dong J, Liu J, Bigio EH, Mesulam M, Wang T, Sun L, Wang L, Lee AY, McGee WA, Chen X, Fushimi K, Zhu L, Wu JY. 2019. TDP-43 induces mitochondrial damage and activates the mitochondrial unfolded protein response. PLoS Genetics 15:e1007947. DOI: 10.1371/journal.pgen.1007947.

Watts JC, Condello C, Stöhr J, Oehler A, Lee J, DeArmond SJ, Lannfelt L, Ingelsson M, Giles K, Prusiner SB. 2014. Serial propagation of distinct strains of Aβ prions from Alzheimer’s disease patients. Proceedings of the National Academy of Sciences of the United States of America 111:10323–10328. DOI: 10.1073/pnas.1408900111.

Wickner RB, Shewmaker FP, Bateman DA, Edskes HK, Gorkovskiy A, Dayani Y, Bezsonov EE. 2015. Yeast prions: structure, biology, and prion-handling systems. Microbiology and Molecular Biology Reviews 79:1–17. DOI: 10.1128/MMBR.00041-14.

Woerner AC, Frottin F, Hornburg D, Feng LR, Meissner F, Patra M, Tatzelt J, Mann M, Winklhofer KF, Hartl FU, Hipp MS. 2016. Cytoplasmic protein aggregates interfere with nucleocytoplasmic transport of protein and RNA. Science 351:173–176. DOI: 10.1126/science.aad2033.

Yuan AH, Hochschild A. 2017. A bacterial global regulator forms a prion. Science 355:198–201. DOI: 10.1126/science.aai7776.

